# Role of SpoIVA ATPase Motifs During *Clostridioides difficile* Sporulation

**DOI:** 10.1101/2020.07.02.183343

**Authors:** Hector Benito de la Puebla, David Giacalone, Alexei Cooper, Aimee Shen

**Affiliations:** Department of Molecular Biology and Microbiology, Tufts University School of Medicine, Boston, Massachusetts, USA; Graduate Program in Molecular Biology and Microbiology, Tufts University School of Medicine, Boston, Massachusetts, USA; Department of Microbiology and Molecular Genetics, University of Vermont, Burlington, Vermont, USA

## Abstract

The nosocomial pathogen, *Clostridioides difficile*, is a spore-forming obligate anaerobe that depends on its aerotolerant spore form to transmit infections. Functional spore formation depends on the assembly of a proteinaceous layer known as the coat around the developing spore. In *C. difficile*, coat assembly depends on the conserved coat protein, SpoIVA, and the clostridial-specific coat protein, SipL, which directly interact. Mutations that disrupt their interaction cause coat to mislocalize and decrease functional spore formation. In *B. subtilis*, SpoIVA is an ATPase that uses ATP hydrolysis to help drive its polymerization around the forespore. Loss of SpoIVA ATPase activity impairs *B. subtilis* SpoIVA encasement of the forespore and activates a quality control mechanism that eliminates these defective cells. Since this mechanism is lacking in *C. difficile*, we tested whether mutations in *C. difficile*’s SpoIVA ATPase motifs impair functional spore formation. Disrupting *C. difficile* SpoIVA ATPase motifs resulted in phenotypes that were typically >10^4^ less severe than the equivalent mutations in *B. subtilis*. Interestingly, mutation of ATPase motif residues predicted to abrogate SpoIVA binding to ATP decreased SpoIVA-SipL interaction, whereas mutation of ATPase motif residues predicted to disrupt ATP hydrolysis but retain binding to ATP enhanced SpoIVA-SipL interaction. When a *sipL* mutation known to reduce binding to SpoIVA was combined with a *spoIVA* mutation predicted to prevent SpoIVA binding to ATP, spore formation was severely exacerbated. Since this phenotype is allele-specific, our data implies that SipL recognizes the ATP-bound form of SpoIVA and highlights the importance of this interaction for functional *C. difficile* spore formation.

**Importance:** The aerotolerant spores formed by the major nosocomial pathogen *Clostridioides difficile* are its primary infectious particle. However, the mechanism by which this critical cell type is assembled remains poorly characterized, especially with respect to its protective coat layer. We previously showed that binding between the spore morphogenetic proteins, SpoIVA and SipL, regulates coat assembly around the forespore. SpoIVA is widely conserved among spore-forming bacteria, and its ATPase activity is essential for *Bacillus subtilis* to form functional spores. In this study, we determined that mutations in *C. difficile* SpoIVA’s ATPase motifs result in relatively minor defects in spore formation in contrast with *B. subtilis*. Nevertheless, our data suggest that SipL preferentially recognizes the ATP-bound form of SpoIVA and identify a specific residue in SipL’s C-terminal LysM domain that is critical for recognizing the ATP-bound form of SpoIVA. These findings advance our understanding of how SpoIVA-SipL interactions regulate *C. difficile* spore assembly.

## Introduction

The Gram-positive spore-former, *Clostridioides difficile*, formerly known as *Clostridium difficile*, is a leading cause of healthcare-associated infections worldwide (1). *C. difficile* causes a debilitating diarrhea that can lead to severe complications like pseudomembranous colitis and toxic megacolon (2). These disease symptoms are caused by the glucosylating toxins produced by *C. difficile* (3), although disease typically only occurs in individuals experiencing gut dysbiosis because dysbiosis is required for *C. difficile* to establish a replicative niche in the gut (4, 5). While *C. difficile* is intrinsically resistant to many antibiotics, its metabolically dormant spore form further facilitates *C. difficile* survival, since spores are inert to antibiotics and can persist in the environment for long periods of time (6).

Spores are also the major transmissive form of *C. difficile*, since it is an obligate anaerobe (7, 8). As a result, spore formation is critical for *C. difficile* to survive exit from the host and persist in the environment (9). This important developmental process involves a series of coordinated morphological changes that begins with the formation of a polar septum, which creates a larger mother cell and smaller forespore (10, 11). The mother cell then engulfs the forespore, suspending the double-membraned forespore within the cytosol of the mother cell. Following engulfment, a thick layer of modified peptidoglycan known as the cortex forms between the forespore’s two membranes. The cortex layer is critical for maintaining metabolic dormancy and confers resistance to heat and ethanol (12). As the forespore develops, a series of proteins localizes to and assembles on the outer forespore membrane to form concentric proteinaceous shells around the forespore known as the coat, which protects spores against enzymatic and chemical insults like lysozyme and quaternary amines (13).

The molecular mechanisms controlling these developmental stages have been extensively characterized in *Bacillus subtilis*, and factors critical for each of these morphological stages have been identified (10). While most of these *B. subtilis* morphogenetic factors are conserved across spore-formers (14, 15), analyses of spore assembly in *C. difficile* indicate that many of these factors have different functions or requirements in *C. difficile*. The conserved transmembrane protein, SpoIIM, is essential for engulfment in *B. subtilis* (16) but dispensable for this process in *C. difficile* (17, 18), while the lytic transglycosylase, SpoIID, is essential for engulfment in *B. subtilis* (19) but only partially required in *C. difficile* (17, 18). Although components of a channel complex are required to maintain the integrity of the forespore in both *B. subtilis* and *C. difficile* (20-22), this complex is required for engulfment in *C. difficile* (21, 22) yet dispensable in some conditions in *B. subtilis* (23).

In addition, *C. difficile* and *B. subtilis* differ considerably in how they assemble the coat layer, since only two of the nine coat morphogenetic proteins identified in *B. subtilis* have homologs in *C. difficile*: SpoVM and SpoIVA (24). While both these proteins are conserved across spore formers (14, 15), they are differentially required in *C. difficile* relative to *B. subtilis*, since SpoVM is largely dispensable for functional spore formation in *C. difficile* (25) in contrast with *B. subtilis* (26). In *B. subtilis*, SpoIVA and SpoVM are the first proteins to be recruited to the outer forespore membrane during engulfment, and both are essential for coat and cortex assembly and thus functional spore formation (26-30). SpoVM recognizes the positive curvature of the forespore and embeds itself in this membrane (28, 31). SpoVM directly recruits SpoIVA to the forespore (32) and potentiates the polymerization of SpoIVA around the forespore (33). SpoIVA is a cytoskeletal-like protein with ATPase activity that encases the forespore in an ATP-dependent manner (34, 35) when bound to SpoVM (32). SpoIVA subsequently recruits additional coat morphogenetic proteins (27, 36). Thus, these two proteins form the basement layer on which the coat assembles in *B. subtilis*.

While *C. difficile* SpoIVA also regulates coat assembly and is essential for functional spore formation, *C. difficile spoIVA* mutants produce cortex (37) unlike their counterparts in *B. subtilis* (27). Furthermore, *C. difficile spoVM* mutants exhibit only a ∼3-fold decrease in functional, heat-resistant spore formation (25) compared to the ∼10^6^ defect observed in *B. subtilis spoVM* mutants (26). The relatively minor defect of *C. difficile spoVM* mutant spores is likely because *C. difficile* does not encode a quality control pathway that eliminates defective sporulating cells in *B. subtilis* (29, 38).

This quality control mechanism senses defects in *B. subtilis* SpoIVA localization around the forespore, which can occur due to loss of SpoVM or SpoIVA ATPase activity (29). Notably, the quality control pathway only appears to be conserved in spore-formers of the order *Bacilliales* (39), suggesting that clostridial organisms either lack a mechanism for eliminating spores with coat assembly defects or employ a clostridial-specific quality control mechanism.

Another major difference in coat assembly between *C. difficile* and *B. subtilis* is the requirement for SipL, a SpoIVA-interacting coat morphogenetic protein that is only conserved in the Clostridia (37). *C. difficile sipL* mutants phenocopy *C. difficile spoIVA* mutants in that they exhibit similar defects in coat localization and heat-resistant spore formation despite making a cortex layer (37). We previously showed that SipL directly binds SpoIVA through SipL’s C-terminal LysM domain (37) and that disrupting SipL LysM domain binding to SpoIVA via mutation of specific residues causes defects in coat assembly and heat-resistant spore formation (40).

While SipL residues important for recognizing SpoIVA have been identified, the SpoIVA residues recognized by SipL’s LysM domain remain unknown. We previously showed that mutating a residue predicted to prevent *C. difficile* SpoIVA from binding ATP also reduced SpoIVA binding to SipL in co-affinity purification assays performed on *E. coli* lysates (37). Since previous work with *B. subtilis* SpoIVA revealed that ATP binding to SpoIVA induces a conformational change that allows SpoIVA to self-polymerize upon hydrolyzing ATP (34, 41), we sought to test whether ATP binding and/or hydrolysis were required for *C. difficile* SipL to recognize SpoIVA during spore formation. Since the ATPase activity of *B. subtilis* SpoIVA is also required for SpoIVA to fully encase the forespore (34), we wondered how disrupting conserved ATPase motifs in *C. difficile* SpoIVA would impact spore coat assembly. To this end, we mutated *C. difficile* SpoIVA’s conserved ATPase motifs and determined their impact on *C. difficile* spore formation, binding to SipL, and coat localization. Our analyses indicate that SipL preferentially recognizes the ATP-bound form of SpoIVA and reveal further differences in the functional requirements for conserved spore proteins between *B. subtilis* and *C. difficile*.

## Results

### Alanine mutations in SpoIVA ATPase motifs do not strongly reduce *C. difficile* heat-resistant spore formation

The N-terminus of SpoIVA has homology to the translation factor (TRAFAC) clade of GTPases (**Figure S1**, (34)), since SpoIVA carries three of four highly conserved motifs that distinguish TRAFAC GTPases (and the larger P-loop NTPase superfamily) (42). These three motifs include the (i) Walker A/P-loop motif, (ii) Sensor threonine (also known as the switch I motif), and (iii) Walker B/switch II motif. The Walker A motif is required for ATP binding and hydrolysis, while the Sensor threonine and Walker B motifs are required for ATP hydrolysis because they help coordinate the Mg^2+^ ion co-factor. SpoIVA homologs lack the fourth motif, which confers specificity for GTP over ATP (42), consistent with the finding that SpoIVA hydrolyzes ATP instead of GTP (35).

Mutation of the three strictly conserved motifs in *B. subtilis* SpoIVA indicate that they are critical for SpoIVA function during sporulation (34, 35). Substitution of the conserved lysine in the Walker A motif to glutamate (K30E) prevented ATP binding *in vitro* and decreased *B. subtilis* heat-resistant spore formation by 10^8^. However, if the lysine was changed to an alanine (K30A), heat-resistant spore formation decreased only 20-fold relative to wild type (35). The relatively mild defect of a K30A mutation could result from the trace amount of ATP binding observed with SpoIVA_K30A_, which binds ATP ∼20-fold less efficiently than wild-type SpoIVA (35). Mutation of the strictly conserved Sensor threonine to alanine (T70A) and its neighboring threonine (T71A) decreased heat-resistant spore formation by ∼10^4^ (34). (The dual mutations overcome the ability of Thr71 to partially substitute for the T70A mutation). Finally, mutation of the conserved aspartate (Asp97) in SpoIVA’s Walker B motif reduced heat-resistant spore formation in *B. subtilis* by ∼10^7^ (34). Notably, Sensor threonine and Walker B mutants retain the ability to bind ATP but cannot hydrolyze it (34).

Since all three of these NTPase motifs are conserved in *C. difficile* SpoIVA, we tested whether the Walker A, Sensor threonine, and Walker B motifs were critical for *C. difficile* SpoIVA function during spore formation. We constructed mutations analogous to those generated in *B. subtilis* in these three motifs (34, 35). We mutated lysine 35 in the Walker A motif to either alanine or glutamate (K35A, K35E); Sensor threonine 75 to alanine (T75A); and aspartate 102 in the Walker B motif to alanine (D102A). Since *C. difficile* SpoIVA does not carry a second threonine next to the Sensor threonine, only a single threonine point mutation was made. The mutant alleles were introduced into either (i) the ectopic *pyrE* locus using *spoIVA* complementation constructs (43) or (ii) the native *spoIVA* locus using allelic exchange. We then determined the impact of these mutations on heat-resistant spore formation using a heat resistance assay. This assay measures the ratio of spores relative to total cells in sporulating cultures based on the ability of spores to survive heat treatment and outgrow to form colonies when plated on media containing germinant (44). As a result, decreases in the heat resistance ratio can be caused by defects in spore formation, heat resistance, germination, and/or outgrowth.

Strains encoding alanine substitutions in the Walker A, Sensor threonine, and Walker B motifs (*K35A, T75A*, and *D102A*) resulted in only ∼3-fold reductions in heat-resistant spore formation relative to wild type, regardless of whether the mutant *spoIVA* alleles were expressed from their native locus (**Figure 1**) or the *pyrE* locus (**Figure S2**). Mutation of the Walker A lysine to glutamate (*K35E*), led to more severe defects in heat-resistant spore formation relative to wild type (∼100-fold defect), regardless of the chromosomal location of the *K35E* allele (**Figures 1A** and **S1A**). The phenotype of the *C. difficile K35A* mutant was ∼7-fold less severe than an equivalent mutant in *B. subtilis* (35), while the phenotypes of the Sensor threonine and Walker B mutants in *C. difficile* were at least 4,000-fold less severe than their counterparts in *subtilis* (34). While the heat resistance defect for *K35E* mutant was ∼10^2^ lower than wild-type *C. difficile* (**Figures 1A** and **S1**), the equivalent mutation in *B. subtilis* led to a 10^8^ decrease in heat resistance (35). Thus, mutations in conserved NTPase motifs in *C. difficile* SpoIVA have relatively minor effects relative to the equivalent mutations in *B. subtilis*.

**Figure 1.**
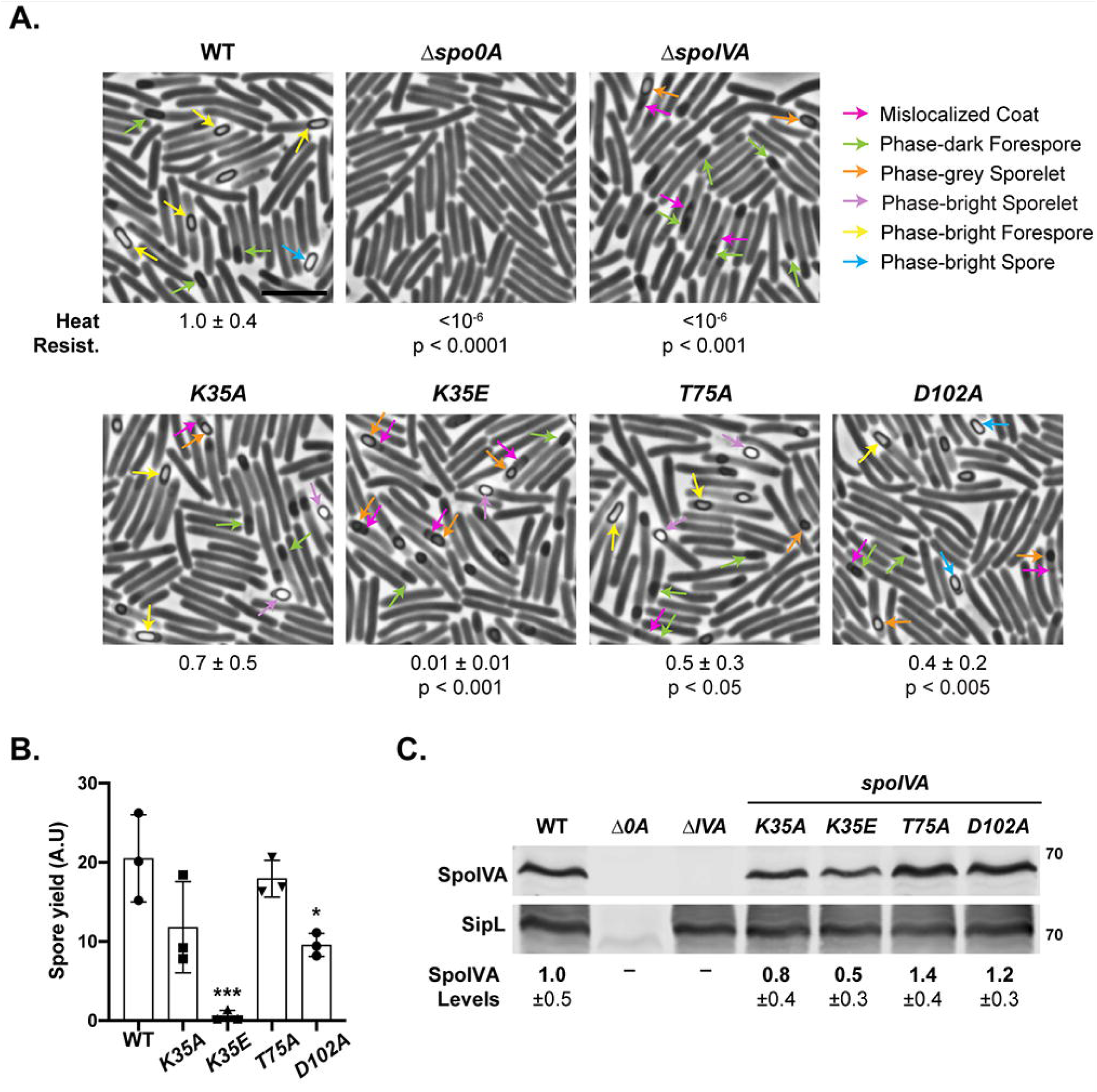
Effect of SpoIVA ATPase motif mutations encoded in native locus on functional spore formation and spore purification efficiency. (A) Phase-contrast microscopy analyses of the indicated *C. difficile* strains ∼20 hrs after sporulation induction. The SpoIVA ATPase motif mutations are encoded in the native *spoIVA* locus. Arrows mark examples of sporulating cells at different stages of maturation: pink arrows mark regions of mislocalized coat based on previous studies (21, 25); green arrows highlight immature phase-dark forespores; orange arrows highlight phase-gray sporelets, which look swollen and are surrounded by a phase-dark ring; purple arrows highlight phase-bright sporelets, which are swollen and surrounded by a phase-dark ring; yellow arrows mark mature phase-bright forespores; phase-brightness reflects cortex formation (39, 56); blue arrows highlight phase-bright free spores. Heat resistance efficiencies are based on 20-24 hr sporulating cultures and represent the mean and standard deviation for a given strain relative to wild type based on a minimum of three biological replicates. Scale bar represents 5 μm. The limit of detection of the assay is 10^−6^. (B) Spore yields based on purifications from the indicated strains from three biological replicates. Yields were determined by measuring the optical density of spore purifications at 600 nm; yields are expressed in arbitrary units. (B) Western blot analyses of SpoIVA and SipL. SpoIVA levels were quantified based on analyses of three biological replicates using (57). Statistical significance for all assays was determined relative to wild type using a one-way ANOVA and Tukey’s test. No statistically significant differences were detected for the western blotting data. *** p < 0.005.

To determine the extent to which the ATPase motif mutations affected the number of spores produced, we measured the efficiency with which spores from the ATPase motif mutant strains could be purified. Spores were harvested from sporulating cultures grown on equal numbers of plates and purified using a Histodenz gradient (**Figure 1B**). While the purification yield for the *T75A* mutant spores did not change relative to wild type, the purification yields for *K35A* and *D102A* mutant spores decreased by ∼2-fold, although only the decrease for *D102A* mutant spores was statistically significant (p < 0.05). The purification yield for *K35E* mutant spore was ∼50-fold (p < 0.001) that of wild type.

Phase-bright spores resembling wild-type spores were purified from the ATPase motif mutants carrying alanine mutations, although circular spores and spores with appendages were more frequently observed in the mutant strains relative to wild-type (**Figure S3**). The small amount of *K35E* spores that could be purified were primarily phase-bright sporelets, which are smaller and more swollen than wild type spores (25, 40). Taken together, our results suggest that the *K35E* mutant’s heat resistance defect is mainly caused by a defect in spore assembly.

### ATPase motif mutations do not strongly affect SpoIVA levels in sporulating cells

Since we previously found that mutating the Walker A lysine to glutamate reduced SpoIVA_K35E_ levels when recombinantly produced in *E. coli* (37), we measured the levels of SpoIVA and its binding partner SipL in the *C. difficile* ATPase motif mutant strains using Western blotting. These analyses revealed that SpoIVA levels were slightly but consistently reduced (∼2-fold) in the *K35E* mutant, although the difference was not statistically significant (**Figure 1C**). The decrease in SpoIVA_K35E_ suggests that the K35E point mutation partially destabilizes SpoIVA in *C. difficile*, consistent with analyses in *E. coli* (37). However, it is unclear whether the ∼2-fold decrease in SpoIVA_K35E_ protein levels contributes significantly to the ∼100-fold heat resistance defect of the *K35E* mutant.

### Mutation of SpoIVA’s ATPase motifs leads to morphological defects in spore formation

Even though the effects of mutating *C. difficile* SpoIVA’s ATPase motifs to alanine were relatively minor in the heat resistance assay, visual inspection of the purified spores (**Figure S3**) and sporulating cells (**Figures 1** and **S2**) by phase-contrast microscopy revealed that the mutant spores had morphological defects relative to wild type. K35A, T75A, and D102A mutants often produced sporelets, which are brighter and more swollen than the oblong forespore typically observed in wild type (18). The alanine mutants also appeared to have mislocalized coat associated with the forespore, since phase-grey areas appeared to extend like a “beard” from the forespore, a phenotype we previously observed in *spoVM* mutant cells (25). The *K35E* mutant produced phase-dark forespores, phase-dark sporelets, and phase-bright sporelets. Mislocalized coat was also observed associated with *K35E* mutant forespores, with the “beards” appearing to extend further into the mother cell cytosol of the *K35E* mutant and the parental Δ*spoIVA* strain (pink arrows, **Figures 1A** and **S2A**).

To visualize mislocalized coat with greater resolution, we analyzed the ATPase motif mutants using transmission electron microscopy (TEM). Analyses of ∼50 sporulating cells that had completed engulfment revealed that Walker A motif mutants frequently exhibited coat encasement defects, with the coat encasing the forespore in only ∼20% of *K35A* mutant spores and ∼5% of *K35E* mutant spores (**Figure 2**). The coat encasement defect of the *K35E* mutant was almost as severe as Δ*spoIVA*, for which no coat encasement was observed.

**Figure 2.**
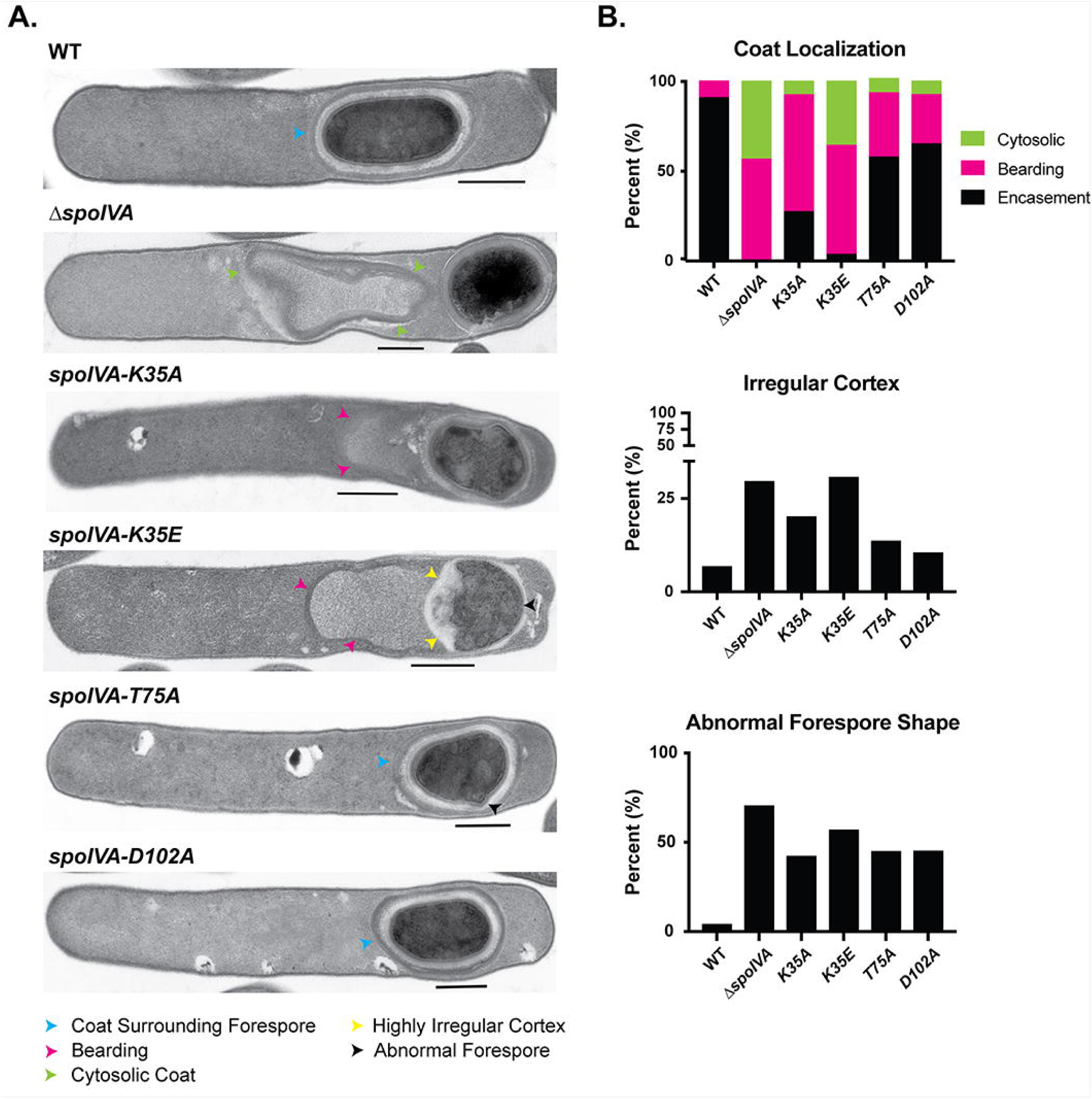
Coat and cortex abnormalities in *spoIVA* ATPase motif mutants. (A) Transmission electron microscopy (TEM) analyses of wild type, Δ*spoIVA*, and *spoIVA* mutants encoding ATPase motif mutations in their native locus 23 hrs after sporulation induction. Scale bars represent 500 nm. Blue arrows mark properly localized coat, i.e. surrounding the entire forespore (FS); yellow arrows mark where coat appears to detach from the forespore but stays partially associated, also known as “bearding” (25); and green arrows mark where the coat has fully detached from the forespore and is entirely cytosolic. A forespore with a cortex layer that varies markedly in thickness is highlighted with a yellow arrow. A misshapen forespore is marked with a black arrow. At least 50 cells for each strain with visible signs of sporulation from a single biological replicate were analyzed.

The Walker A mutations resulted in coat appearing to slough off the forespore (termed “bearding”) at a frequency similar to the parental Δ*spoIVA* sporulating cells (50-60%, **Figure 2**). We previously termed this phenotype “bearding” (25). The *K35E* mutation also increased the frequency of the coat completely mislocalizing to the cytosol similar to the Δ*spoIVA* mutant (∼35-40% of cells). Coat mislocalization defects were comparatively less severe in the Sensor threonine and Walker B mutants, *T75A* and *D102A*, with the majority of these mutants completing coat encasement. Nevertheless, bearding was still observed in ∼20-30% of these mutant cells (**Figure 2**).

In addition to the coat localization defects, SpoIVA ATPase motif mutants were more likely to produce cortex layers of highly irregular thickness than wild-type cells. Over 30% of *K35E* and Δ*spoIVA* sporulating cells produced highly irregular cortex layers (**Figures 2**), while this defect was observed in only ∼5% of wild-type cells. Irregular cortex was also observed in 20% of *K35A* and ∼10% of *T75A* and *D102A* mutants. These cortex abnormalities were frequently associated with changes in forespore shape. Notably, almost half of the alanine ATPase motif mutants and even more of the *K35E* and Δ*spoIVA* mutants produced forespores that were irregularly shaped, whether they were more circular in shape (as seen by phase-contrast microscopy, (**Figure 1**), exhibited protrusions, or appeared wavy (**Figure 2**). Since we have reported a similar defect in cortex morphology in *spoVM* mutants (25), SpoVM and SpoIVA would appear to modulate cortex synthesis on an unknown level in *C. difficile*.

### ATPase motif mutations decrease SpoIVA encasement of the forespore

We next sought to determine the effects of the ATPase motif mutations on SpoIVA localization around the forespore because *B. subtilis* mutants lacking ATPase activity fail to fully encase the forespore (34, 35). In *B. subtilis*, disrupting ATP binding by mutating the Walker A lysine to alanine (*K30A*) caused SpoIVA to localize to a single cap at the mother cell proximal pole, while mutation of the lysine to glutamate (*K30E*) redistributed SpoIVA to the cytosol (35). Disrupting ATP hydrolysis (but not ATP binding) due to mutations of the Walker B motif in *B. subtilis* still allowed SpoIVA to localize to both poles of the forespore but prevented full encasement of the forespore in >80% of cells (34).

To assess whether the *C. difficile* ATPase motif mutations decreased the ability of SpoIVA to encase the forespore, we introduced *mCherry-spoIVA* alleles carrying the ATPase motif mutations into the ectopic *pyrE* locus of strains encoding the same *spoIVA* mutant allele in their native locus (e.g. *K35A*/*mCherry-spoIVA*_*K35A*_). These strains encode an untagged SpoIVA ATPase motif mutant along with an mCherry-SpoIVA ATPase motif mutant because wild-type mCherry-SpoIVA is not fully functional unless it is co-produced with untagged wild-type SpoIVA (25).

While wildtype mCherry-SpoIVA localized primarily around the forespore in ∼80% of cells ∼24 hrs after sporulation was induced, mutation of the ATPase motifs reduced mCherry-SpoIVA encasement of the forespore (**Figure 3**). The Sensor threonine and Walker B mutants encased the forespore ∼2-fold lower frequency relative to wild type, with T75A and D102A mCherry-IVA either partially encasing the forespore or capping both poles of the forespore. The localization defects were slightly more severe in the *K35A* Walker A motif mutant relative to the *T75A* and *D102A* alanine mutants. In contrast, the K35E mutation resulted in mCherry-SpoIVA mainly exhibiting a capped distribution (either single or double capped) and surrounding the forespore in only ∼20% of cells.

**Figure 3.**
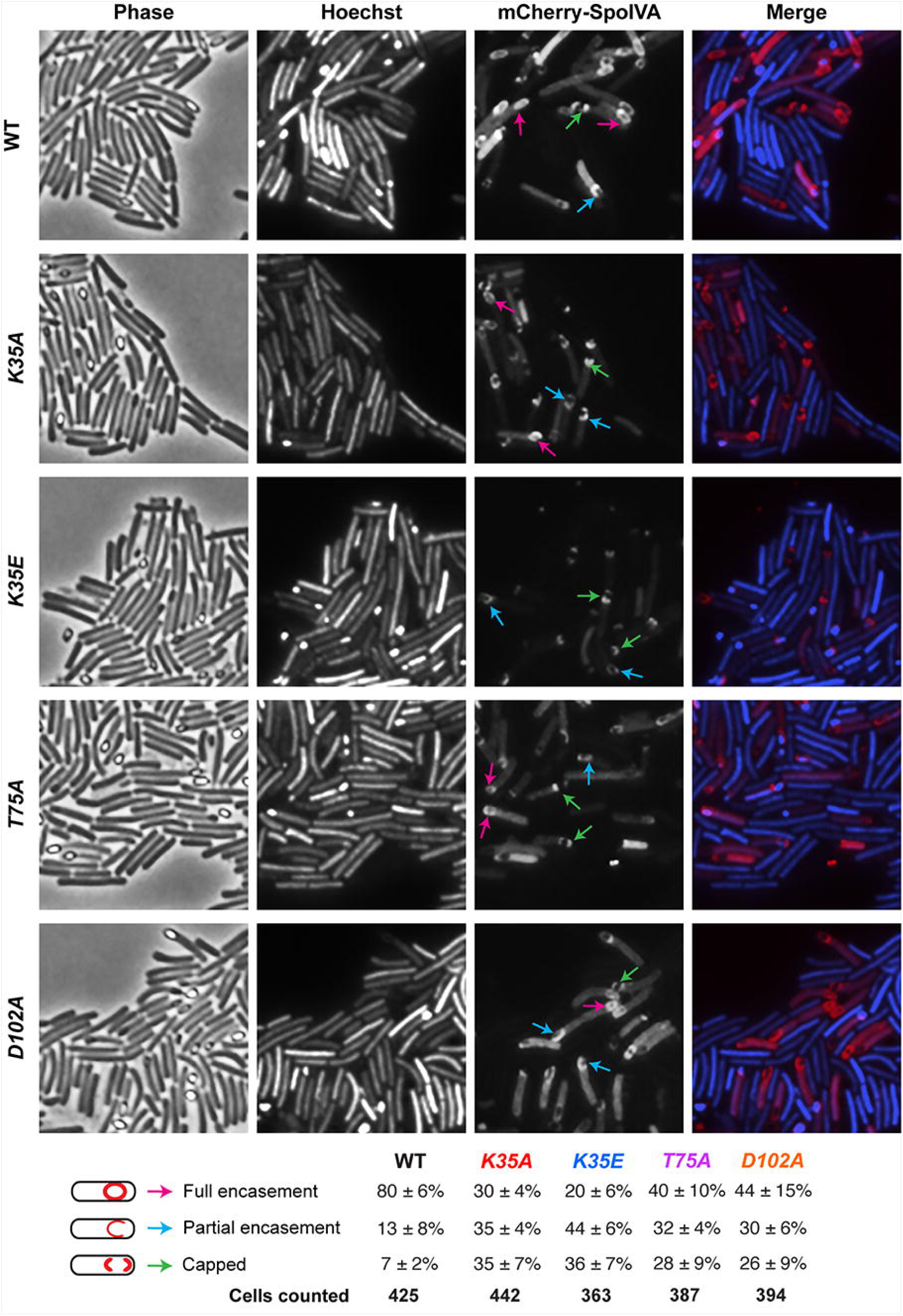
Effect of SpoIVA ATPase motif mutations on SpoIVA localization. Fluorescence microscopy analyses of wild type and *spoIVA* mutants encoding ATPase motif mutations in their native locus 23 hrs after sporulation induction. Phase-contrast microscopy was used to visualize sporulating cells (Phase). Hoechst staining used to visualize the nucleoid is shown in blue, and mCherry-SpoIVA fluorescence is shown in red. The merge of the Hoechst staining and mCherry signal is shown. Schematic of coat localization phenotypes quantified is shown along with the percentage of cells in a given strain that exhibited this phenotype. The average percentages and standard deviations shown are based on counts from three biological replicates, with multiple images from each replicate being quantified. The total number of cells counted is also shown.

The K35E mutation noticeably diminished the intensity of mCherry-IVA signal relative to wild type and the other ATPase motif mutants, but this was not due to mCherry being liberated from mCherry-SpoIVA_K35E_ through proteolysis (**Figure S4**). Less mCherry-IVA_K35E_ was observed by western blotting, however, consistent with western blot analyses of the untagged SpoIVA variant (**Figure 1**). Taken together, mutating *C. difficile* SpoIVA ATPase motifs decreases SpoIVA encasement around the forespore but not as severely as in *B. subtilis* (34, 35).

### Coat protein encasement of the forespore is reduced in SpoIVA ATPase motif mutants

*C. difficile* SpoIVA is required to recruit SipL to the forespore (40), so we tested whether the ATPase motif mutations decreased SipL localization using a SipL-mCherry fluorescent protein fusion we previously constructed (40). Since this fusion protein encases the forespore more efficiently when SipL-mCherry is the only form of SipL present in the cell (40), we deleted the *sipL* gene from the *spoIVA* mutants encoding ATPase motif mutations in their native loci. We then integrated a construct encoding SipL-mCherry into the *pyrE* locus. While SipL-mCherry localized to the forespore in all the ATPase motif mutants tested, SipL-mCherry encased the forespore ∼2-3-fold less frequently in these mutants relative to wild type (**Figure 4**). Indeed, in most (>70%) SpoIVA ATPase motif mutant cells, SipL-mCherry was detected “capping” either both poles of the forespore or the pole closest to the mother cell.

**Figure 4.**
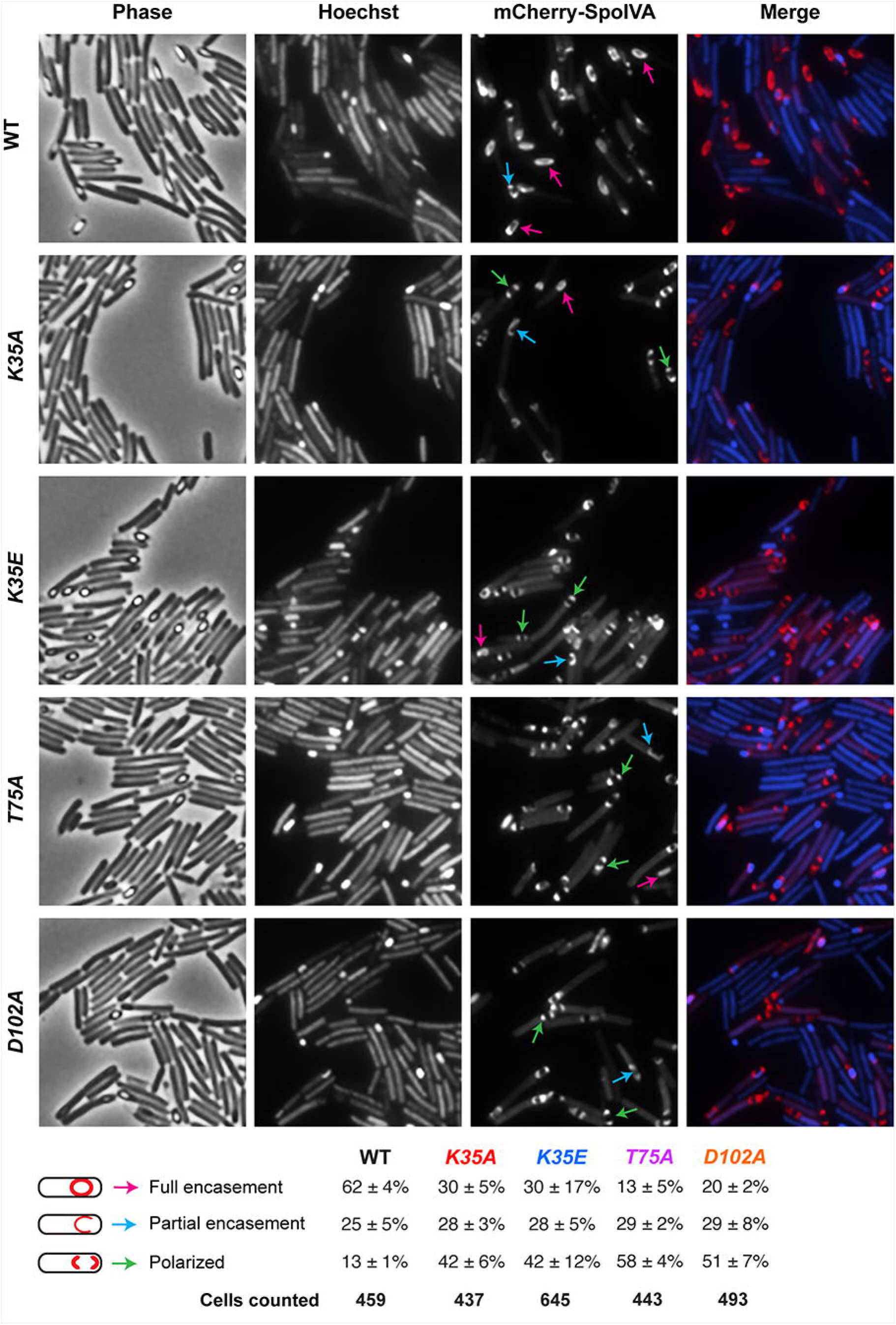
Effect of SpoIVA ATPase motif mutations on SipL localization. Fluorescence microscopy analyses of Δ*sipL* complemented with *sipL-mCherry* (designated “WT”) and *spoIVA* mutants encoding ATPase motif mutations in their native locus from which *sipL* was deleted and *sipL-mCherry* was integrated into the *pyrE* locus. Microscopy was performed on samples 23 hrs after sporulation induction. Phase-contrast microscopy was used to visualize sporulating cells (Phase). Hoechst staining used to visualize the nucleoid is shown in blue, and SipL-mCherry fluorescence is shown in red. The merge of the Hoechst staining and mCherry signal is shown. Schematic of coat localization phenotypes quantified is shown along with the percentage of cells in a given strain that exhibited this phenotype. The average percentages and standard deviations shown are based on counts from three biological replicates, with multiple images from each replicate being quantified. The total number of cells counted is also shown.

To test whether the SpoIVA ATPase motif mutations affected the localization of coat proteins found in the outer layers of the spore as suggested by the coat bearding observed by TEM (**Figure 2**), we analyzed the localization of an mCherry fusion to the spore surface protein, the CotE mucinase (45). To localize *C. difficile* CotE, we introduced a construct encoding a CotE-mCherry fusion protein (18) into the *pyrE* locus of the ATPase motif mutant strains. In wild-type cells, *C. difficile* CotE encases the forespore in a typically non-uniform fashion, concentrating at the forespore poles as two caps (18, 21). In the absence of SpoIVA, CotE-mCherry failed to localize to the forespore and instead formed a bright focus in the cytosol of sporulating cells (**Figure S5**), as seen with prior studies of a CotE-SNAP fusion protein in a *spoIVA* mutant (21). However, when the SpoIVA ATPase motifs were mutated to alanine, CotE-mCherry still localized to both poles of the forespore (**Figure S5**), although CotE-mCherry formed a single bright focus close to the forespore of the *K35E* Walker A mutant (rather than two polar caps) more frequently than the other mutant strains. Unfortunately, the non-uniform distributions of CotE-mCherry around the forespore made it difficult to quantify the different localization patterns observed. Regardless, relative to a *spoIVA* null mutant, the *C. difficile* ATPase motif mutations did not severely disrupt the localization of CotE-mCherry.

### Mutation of the SpoIVA Walker A motif decreases SpoIVA binding to SipL

Our localization analyses with SipL-mCherry revealed that SipL localizes to the forespore but encases it less efficiently when SpoIVA’s ATPase motifs are mutated (**Figure 4**). Given that loss of SpoIVA causes SipL-mCherry to redistribute to the cytosol (40), our data suggest that the SpoIVA ATPase motif mutants retain the ability to bind and recruit SipL to the forespore. However, prior co-affinity purification analyses with SpoIVA and SipL in *E. coli* indicated that the SpoIVA K35E strongly impairs binding to SipL (37). To address this potential discrepancy, we quantified the interaction between SpoIVA and SipL using (i) a bacterial two hybrid assay in *E. coli* and (ii) co-immunoprecipitation analyses in *C. difficile* (**Figure 5**).

**Figure 5.**
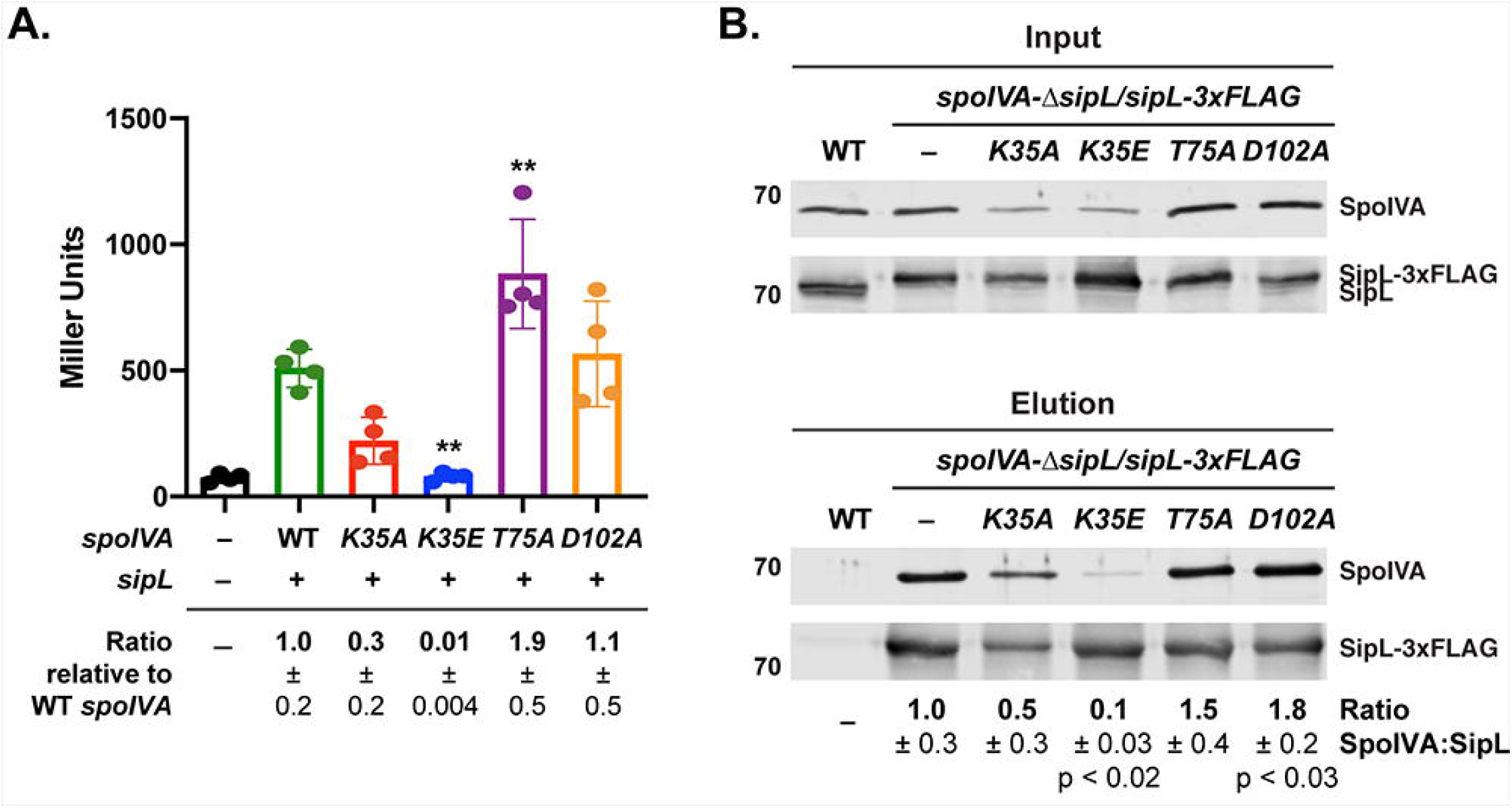
Binding of SipL to SpoIVA ATPase Motif Mutants. (A) Bacterial adenylate cyclase two hybrid (BACTH) analysis of SipL binding to SpoIVA ATPase motif mutants. Plasmids encoding pKNT25-*sipL* and pUT18C-*spoIVA* ATPase motif variants were co-transformed into *E. coli* BTH101. After incubating the transformation plates at 30°C for 40 hrs, cells were scraped off the plate, and β-galactosidase activity (shown in Miller units) was measured to quantify the interaction between SipL and the SpoIVA variants. Assays were performed in technical triplicate for three biological replicates. (–) refers to empty vector. Statistical significance for all assays was determined relative to wild type using a one-way ANOVA and Tukey’s test. ** p < 0.01. (B) Co-immunoprecipitation analysis of SpoIVA ATPase motif mutants binding to FLAG-tagged SipL. SipL-3xFLAG was immunoprecipitated from cleared lysates (“Input” fraction) prepared from either wild type (WT), Δ*sipL/sipL-3xFLAG* complementation strain (–), or Δ*sipL/sipL-3xFLAG* strains encoding SpoIVA ATPase motif mutations in their native locus. Proteins bound to anti-FLAG magnetic beads were eluted using FLAG peptide (“Elution” fraction). Wild type served as a negative control for the anti-FLAG beads. Lysates were prepared from strains induced to sporulate for 22 hrs. The immunoprecipitations shown are representative of three biological replicates. The ratio of SpoIVA:SipL in the co-immunoprecipitations is shown relative to wild-type SpoIVA:SipL. Ratios represent the mean and standard deviation for a given strain relative to wild-type SpoIVA based on three biological replicates.

When an adenylate cyclase-based bacterial two-hybrid assay (46) was used to measure SpoIVA ATPase motif mutant binding to SipL, we observed that the Walker A mutations reduced SpoIVA binding to SipL by ∼3-fold for SpoIVA_K35A_ (p = 0.06), and ∼100-fold for SpoIVA_K35E_ binding (p < 0.003, **Figure 5a**). The latter result mirrors those previously obtained with prior co-affinity purification analyses (37). Surprisingly, the Sensor threonine mutation increased SpoIVA binding to SipL by 2-fold relative to wild-type SpoIVA (*T75A*, p < 0.01), while the Walker B mutation did not significantly affect SpoIVA binding to SipL (**Figure 5a**).

While the two-hybrid assay in *E. coli* allowed us to quantify the direct interaction between SpoIVA and SipL, this assay takes place in the absence of other *C. difficile* sporulation proteins and outside the context of the forespore. Since localization of *B. subtilis* SpoVM and SpoIVA to the forespore cooperatively enhances their encasement of the forespore by increasing their local concentrations (33), we considered the possibility that SpoIVA-SipL binding might be stabilized in the context of the forespore. To determine how the SpoIVA ATPase motif mutations affect binding to SipL during sporulation, we immunoprecipitated FLAG-tagged SipL from a previously constructed strain that expresses *sipL-3xFLAG* from the *pyrE* locus of Δ*sipL* (40). We then measured the amount of untagged SpoIVA ATPase motif mutant variants that were also pulled down. To perform these analyses, we used our double *spoIVA-*Δ*sipL* mutant strains where *sipL* has been deleted from strains encoding SpoIVA ATPase motif mutations in the native *spoIVA* locus. We then complemented this double mutant strain with a construct encoding SipL-3xFLAG integrated into the *pyrE* locus such that the only copy of SipL made is FLAG-tagged.

Immunoprecipitation of SipL-3xFLAG from these strains revealed that mutations of the Walker A, but not Sensor threonine or Walker B, motifs reduced the amount of SpoIVA that co-immunoprecipitated with SipL-3xFLAG. Approximately 3-fold less SpoIVA_K35A_ and 10-fold less SpoIVA_K35E_ (p < 0.02) relative to wild-type SpoIVA co-immunoprecipitated with SipL-3xFLAG. While slightly less SpoIVA_K35E_ was observed in the input fraction, unbound SpoIVA_K35E_ was observed in the flow-through fraction, indicating that SpoIVA_K35E_ levels were not limiting in the co-immunoprecipitation analyses (**Figure S6**). The Walker B mutations resulted in more SpoIVA being pulled-down with SipL-3XFLAG, with 1.5-fold more SpoIVA_T75A_ and 1.8-fold more SpoIVA_D102A_ (p < 0.03) being co-immunoprecipitated. Taken together, the two hybrid analyses in *E. coli* and co-immunoprecipitation analyses in sporulating *C. difficile* cells indicate that mutations in the Walker A motif decrease *C. difficile* SpoIVA binding to SipL. They further suggest that SpoIVA Sensor threonine and Walker B mutations enhance SipL binding. Given that Sensor threonine and Walker B mutations have been shown to “trap” *B. subtilis* SpoIVA in an ATP-bound conformation (34), our results imply that SipL preferentially recognizes the ATP-bound form of SpoIVA.

### Synergistic effects of combining SipL and SpoIVA Walker A mutations on functional spore formation

Unfortunately, we could not directly test this hypothesis because *C. difficile* SpoIVA is largely insoluble in *E. coli* unless co-produced with SipL (37), making it difficult to purify sufficient quantities for biochemical analyses. Given that *C. difficile* SpoIVA shares 71% homology with *B. subtilis* SpoIVA, it is likely that the Walker A motif mutations disrupt *C. difficile* SpoIVA ATP binding, whereas the Sensor threonine or Walker B mutations do not. Nevertheless, to examine this hypothesis further, we reasoned that combining the SpoIVA K35A Walker A motif mutation with SipL mutations that reduce binding to SpoIVA might strongly impair the interaction between SpoIVA and SipL binding and exacerbate the relatively minor heat resistance defect (∼3-fold) of the *spoIVA K35A* mutant (**Figure 1**). In contrast, combining SpoIVA’s Sensor threonine or Walker B motif mutations with the same SipL mutations would not be expected to affect heat-resistant spore formation as strongly because the SpoIVA variants presumably retain binding to ATP and thus bind SipL at levels equal or greater to wild-type SpoIVA (**Figure 5**).

We previously identified two SipL mutations that reduce SipL-SpoIVA binding in co-immunoprecipitation analyses and SipL function in heat resistance assays: I463R and W475E (40). The I463R mutation strongly impaired SpoIVA binding to SipL but surprisingly only decreased functional spore formation by ∼10-fold, while the W475E mutation partially reduced SpoIVA binding to SipL but decreased functional spore formation by ∼100-fold (40). Since the W475E mutation appeared to affect SipL function beyond its ability to disrupt binding to SpoIVA, we reasoned that combining the *spoIVA K35A* Walker A allele with the *sipL W475E* allele might impair functional spore formation less severely than combining the *spoIVA* Walker A mutant allele with the *sipL I463R* allele, since the latter allele reduces binding to SpoIVA more severely than the *sipL W475E* allele.

To generate the double point mutant strains, we complemented the *spoIVA-*Δ*sipL* ATPase motif double mutants used in **Figure 4** with *sipL* complementation constructs encoding either the I463R or W475E mutations. We only analyzed the alanine mutations of the SpoIVA ATPase motifs because the relatively minor heat resistance defects of these mutants (2-3-fold, **Figure 1**) would allow us to detect synergistic effects more readily. Combining the *sipL I463R* allele with the *spoIVA K35A* Walker A mutant allele exacerbated the heat resistance defects of the individual mutations so severely that no heat-resistant spores were detected in the *spoIVA*_*K35A*_ *sipL*_*I463R*_ double mutant strain (**Figure 6A**). This represents a 10^4^ decrease in heat resistance relative to the *sipL I463R* single mutant, whose heat resistance defect is slightly more severe than the *spoIVA K35A* single mutant. Consistent with the severe heat resistance defect of the *spoIVA*_*K35A*_ *sipL*_*I463R*_ double mutant, its sporulating cells resembled those of the *sipL* deletion strain (**Figure S7**). In contrast, combining the *sipL I463R* allele with the *spoIVA T75A* Sensor threonine (*spoIVA*_*K35A*_ *sipL*_*I463R*_) resulted in only a 50-fold decrease in heat-resistant spore formation, which is only 6-fold more severe than the single *sipL I463R* mutant. Combining the *sipL I463R* allele with the *spoIVA D102A* Walker A mutant allele resulted in a 1000-fold decrease in heat resistance for the double mutant; this defect was ∼60-fold more severe than the *sipL I463R* single mutant. Interestingly, the severity of the *sipL*_*I463R*_ *spoIVA* ATPase alanine motif double mutants mirrored the binding (affinities) observed in the bacterial two hybrid assay (**Figure 5A**), and the SpoIVA levels in these mutants (**Figure 6B**). SpoIVA_K35A_ was barely detectable in the *spoIVA*_*K35A*_ *sipL*_*I463R*_ double even though its levels were only slightly decreased in the *spoIVA*_*K35A*_ single mutant (**Figure 6B**).

**Figure 6.**
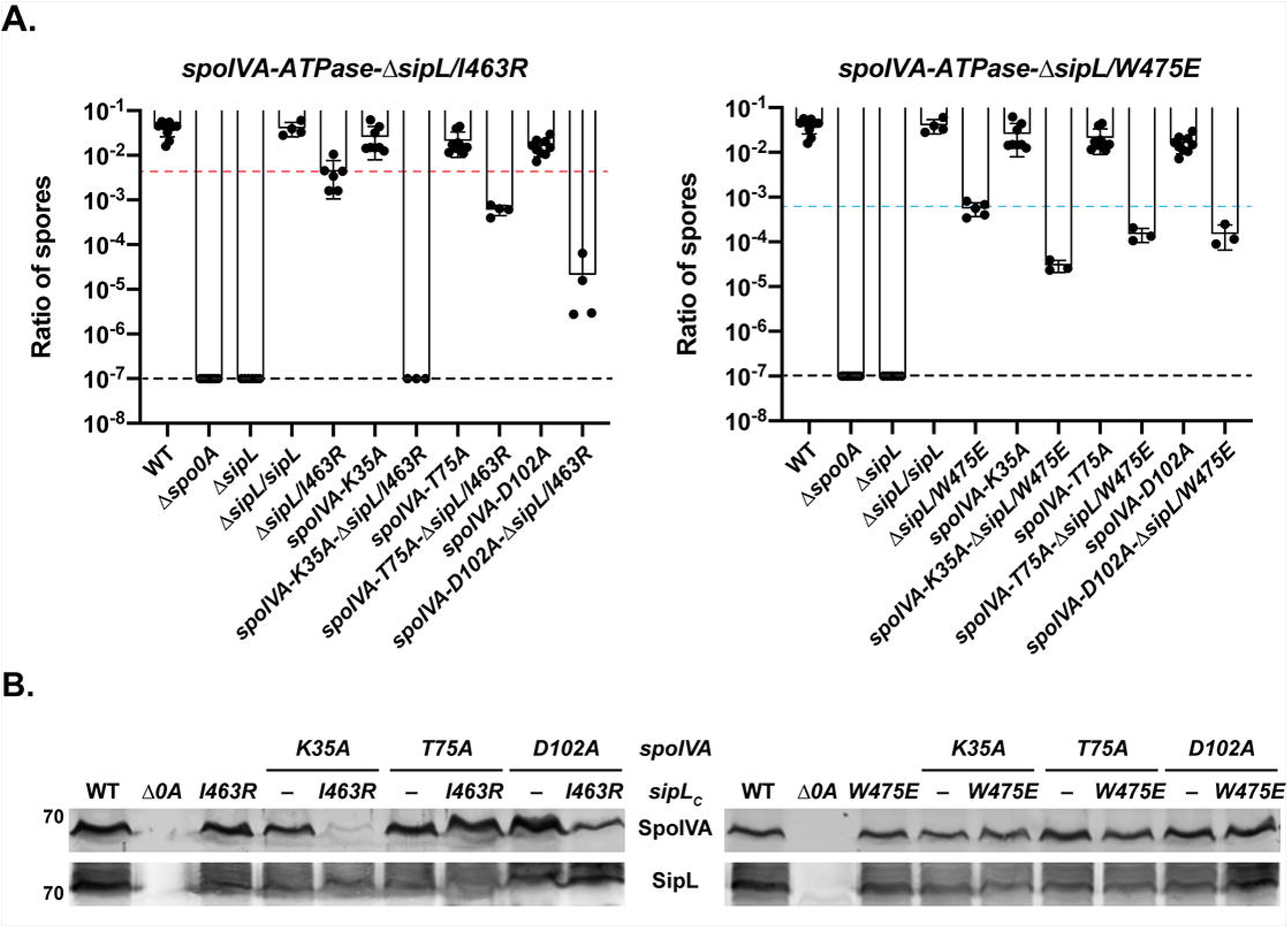
Allele-specific effects of combining SipL and SpoIVA mutations on functional spore formation. (A) The indicated *C. difficile* strains were induced to sporulate for 20-24 hrs. The SpoIVA ATPase motif mutations are encoded in the native *spoIVA* locus, while the SipL LysM domain mutations are encoded in a *sipL* gene integrated into the *pyrE* locus of a Δ*sipL* strain. The ratio of heat-resistant spores to total colony-forming units was determined from a minimum of three biological replicates. The ratio shown is the mean and standard deviation for a given strain relative to wild type. The limit of detection of the assay is 10^−6^ (dotted black line). The dotted red and blue lines indicate the spore ratio for the *sipL I463R* and *W475E* complementation strains, respectively, which had the most severe phenotype of the *sipL* and *spoIVA* single point mutations. Statistical significance for all assays was determined relative to wild type using a one-way ANOVA and Tukey’s test. (B) Western blot analyses of SpoIVA and SipL.

Importantly, when the SpoIVA ATPase motif alanine mutations were combined with the *sipL W475E* allele, the heat resistance defects of the double mutants were largely indistinguishable from each other, and no change in SpoIVA levels was observed in the double mutant strains relative to wild type or the single mutant strains. Since the deleterious effect of combining the Walker A K35A mutation with the SipL I463R mutation appears to be allele-specific, our results are consistent with the hypothesis that SipL specifically recognizes the ATP-bound form of SpoIVA and strongly suggest that Ile463 of SipL plays a critical role in recognizing this form of SpoIVA.

## Discussion

While *B. subtilis* and *C. difficile* use different pathways for localizing coat proteins to the forespore (11, 37, 47), recent work has shown that these organisms can exhibit differential requirements for conserved proteins. In this study, we identify another differential requirement for a conserved spore morphogenetic protein between *B. subtilis* and *C. difficile*. Even though both organisms require the conserved coat morphogenetic protein, SpoIVA, to recruit coat proteins to the forespore and make functional spores (27, 36, 37, 40), we show here that the predicted ATPase activity of SpoIVA is largely dispensable for heat-resistant spore assembly in *C. difficile* in contrast with *B. subtilis*. While alanine mutations that specifically disrupt SpoIVA ATP hydrolysis in *B. subtilis* (i.e. Sensor threonine and Walker B mutations) result in severe (>10^4^) decreases in functional spore formation (34), we determined that the analogous mutations in *C. difficile* SpoIVA result in only 2-3-fold decreases in heat-resistant spore formation (**Figures 1** and **S1**) and minimally disrupt coat localization to the forespore (**Figures 2** and **S5**). Even the most severe ATPase motif mutation we identified, *K35E*, decreased functional *C. difficile* spore formation by only 10^2^ (**Figure 1**), whereas the equivalent mutation reduced spore formation 10^8^ defect in *B. subtilis* (35).

The differential requirement for SpoIVA’s ATPase motifs between *B. subtilis* and *C. difficile* likely reflects the absence of a quality control mechanism for removing spores with defects in encasing SpoIVA around the forespore in *C. difficile*. In *B. subtilis*, this mechanism removes spores with mutations in *spoVM* or *spoIVA*, including SpoIVA ATPase mutants (29), by inducing lysis of the mother cell through the action of a small, *Bacilliales*-specific protein (39), CmpA. The *B. subtilis* quality control mechanism has been proposed to prevent mutations that generate spores of inferior quality from overtaking a population (29, 38). Since *C. difficile spoIVA* ATPase motif mutants and *spoVM* mutants both exhibit coat and cortex abnormalities (**Figures 2, 4**, and **S5**), *C. difficile* would appear to tolerate spores with morphological abnormalities more readily than *B. subtilis*. Given that resistant spore formation is essential for *C. difficile* to survive outside the host and during passage through the stomach (7), the infection cycle presumably places sufficient selective pressure on *C. difficile* to prevent mutants with even minor spore assembly defects from overtaking the population. It would be interesting to test this model by analyzing the effect of the ATPase motif mutations on infectious dose and transmission over many cycles in an animal model of infection, although such analyses would be quite labor intensive and costly.

The differential requirement may also reflect functional redundancy in *C. difficile*’s spore assembly process such that loss of SpoIVA ATPase activity could be compensated by other morphogenetic factors. In *B. subtilis*, SpoVM and SpoIVA form a mutually dependent ratchet that drives assembly of the coat on the forespore surface (33). SpoVM helps recruit SpoIVA to the positively-curved surface of the forespore, while local polymerization of SpoIVA on top of SpoVM enhances SpoVM binding to the forespore membrane (33). Since our data imply that *C. difficile* SipL preferentially binds SpoIVA in its ATP-bound conformation, it is possible that reduced binding of SpoIVA Walker A mutants to SipL is compensated by SpoIVA Walker A mutant binding to SpoVM. For example, the *K35E C. difficile* SpoIVA Walker A mutation decreases binding to SipL by 100-fold in a heterologous bacterium but only 10-fold in co-immunoprecipitation analyses performed in sporulating *C. difficile* (**Figure 5a**). Despite this reduced binding, mCherry-SpoIVA_K35E_ still encases the forespore in 20% of *spoIVA K35E* cells, and SipL-mCherry also encases the forespore of ∼30% of *spoIVA K35E* cells (**Figure 3**). These results suggest that, despite the predicted inability of *C. difficile* SpoIVA_K35E_ to polymerize, other factors (potentially SpoVM) appear to allow *C. difficile* SpoIVA_K35E_ to associate with and encase the forespore even though SpoIVA_K35E_ binds SipL much less efficiently (**Figure 5**). Constructing a *spoVM*-*spoIVA K35E* double mutant would provide insight into this question as would detailed analyses of purified SpoVM, SpoIVA, and SipL binding to membranes similar to the work done with purified *B. subtilis* SpoVM and SpoIVA by Peluso *et al*. (33).

Consistent with the hypothesis that redundant mechanisms stabilize *C. difficile* SpoIVA ATPase motif mutant binding to and encasement of the forespore, *C. difficile spoIVA* ATPase motif alanine mutants fully encased the forespore in ∼30% of Walker A *K35A* mutant cells and ∼40% of Sensor threonine *T75A* and Walker B *D102A* mutants (**Figure 3**). In contrast, the equivalent *B. subtilis* Walker A mutation completely abrogated SpoIVA_K35A_ encasement of the forespore, with SpoIVA_K35A_ only forming a single cap on the mother cell proximal forespore membrane (35), and Sensor threonine and Walker B mutations resulted in only ∼15% of *B. subtilis* fully encasing the forespore (34). While the forespore encasement defects of *B. subtilis* SpoIVA ATPase mutants are consistent with their inability to polymerize into SpoIVA filaments (34, 35), lateral interactions between polymerization-defective SpoIVA proteins may allow mutant SpoIVA to partially encase the forespore (33). These lateral interactions may form more readily with *C. difficile* SpoIVA irrespective of its putative ATPase activity compared to *B. subtilis* SpoIVA given the relatively high percentage of *C. difficile spoIVA* ATPase motif mutant cells that complete forespore encasement. Clearly, an important goal of future work is to determine whether *C. difficile* SpoIVA binds ATP and hydrolyzes it, similar to its *B. subtilis* homolog, for which is shares 50% identity and 69% similarity.

While biochemical analyses of *C. difficile* SpoIVA are challenging because of its low solubility when produced in the absence of SipL in *E. coli* (37), future work should also assess whether *C. difficile* SpoIVA undergoes conformational rearrangements upon binding ATP and then hydrolyzing ATP similar to those reported for *B. subtilis* SpoIVA (34, 41). Determining how *C. difficile* SpoIVA binding to SipL affects these conformational rearrangements will provide important insight into how SpoIVA and SipL collectively regulate spore coat assembly. In addition, crystallographic analyses comparing *C. difficile* SpoIVA alone with SpoIVA bound to SipL would address this question as well as the role of SipL Ile463 in potentially recognizing the ATP-bound form of SpoIVA based on our analyses. Regardless, given the importance of spores to *C. difficile* infection, understanding the mechanisms underlying SpoIVA and SipL function during sporulation could aid in the development of anti-sporulation therapies, which were recently shown to prevent disease recurrence in an animal model of infection (48).

## Materials and Methods

### Bacterial strains and growth conditions

The *C. difficile* strains used are listed in Table S1 (Supplementary materials). All strains derive from the erythromycin-sensitive *630*Δ*erm*Δ*pyrE* parental strain, which is a sequenced clinical isolate 630, which we used for *pyrE-*based allele coupled exchange (ACE) (43). Strains were grown on BHIS, brain heart infusion supplemented with yeast extract and cysteine (49) and taurocholate (TA, 0.1%, [wt/vol]; 1.9 mM), cefoxitin (8 μg/mL) and kanamycin (50 μg/mL) as needed. For ACE, the *C. difficile* defined medium (50) (CDDM) was supplemented with 5-fluorootic acid (5-FOA at [2 mg/mL]) and uracil (at [5 μg/mL]).

The *Escherichia coli* strains used for HB101/pRK24-based conjugations and BACTH assay plasmid preparations are listed in Table S2. *E. coli* strains were grown at 37°C, shaking at 225 rpm in Luria-Bertani (LB) broth. The medium was supplemented with ampicillin (50 μg/mL), chloramphenicol (20 μg/mL) or kanamycin (30 μg/mL) as needed.

### *E. coli* strain construction

All primers used for cloning are listed in Table S2. Details of *E. coli* strain construction are provided in the Supplementary Text S1. All plasmid constructs were sequenced confirmed using Genewiz and transformed into DH5α. HB101/pRK24 *E. coli* strain was used to conjugate sequence-confirmed plasmids into *C. difficile*.

### *C. difficile* strain construction and complementation

Allele-couple exchange (ACE) (43) was used to introduce the *spoIVA* ATPase motif point mutations back into the native locus of a parental Δ*spoIVA*Δ*pyrE* strain. Using this strain facilitated colony PCR screening to identify strains that had restored the native *spoIVA* locus with the mutant gene supplied. ACE was also used to introduce the *sipL* deletion using strain #1704 pMTL-YN3 Δ*sipL* into 630Δ*erm*Δ*pyrE spoIVA* ATPase mutants. Complementations were performed as previously described by conjugating HB101/ pRK24 carrying pMTL-YN1C plasmids into Δ*pyrE-*based strains (51) using allele-coupled exchange.

### Plate-based sporulation assay

*C. difficile* strains were grown overnight from glycerol stocks on BHIS plates supplemented with TA (0.1%, wt/vol). Colonies from these plates were inoculated into BHIS liquid media and back-diluted 1:25 once they were in stationary phase (∼3 hours later). The cultures were grown until they reached an optical density (OD_600nm_) between 0.4 - 0.7; 120 μL was used to inoculate 70:30 agar plates (37) where sporulation was induced for 20 - 24 hours. Cells were analyzed by phase-contrast microscopy and harvested for Western-blot analysis.

### Heat resistance assay on sporulating cells

Heat-resistant spore formation was measured in *C*. difficile sporulating cultures between 20 and 24 hours as previously described (44). Heat-resistance efficiencies represent the average ratio of heat-resistant colony forming units obtained from functional spores in the sample for a given strain relative to the average ratio determined for the wild type based on a minimum of three biological replicates. The ratio of functional, heat-resistant spores shown for a given strain represents the average of every assay performed for this manuscript. Statistical significance was determined using one-way ANOVA and Tukey’s test.

### Western-blot analyses

Samples for Western-blotting analyses were prepared as previously described (37). Briefly, sporulating cell pellets were resuspended in 100 μL of PBS, and 50 μL samples were freeze-thawed for three cycles and then resuspended in 100 μL of EBB buffer (8 M urea, 2 M thiourea, 4% (wt/vol) SDS, 2% (vol/vol) β-mercaptoethanol). The samples were boiled for 20 min, pelleted at high-speed and resuspended in the same buffer to maximize protein solubilization. Finally, the samples were boiled for 5 minutes and pelleted again at high-speed. Samples were resolved on 12% SDS-PAGE gels and then transferred to an Immobilon-FL polyvinylidene difluoride (PVDF) membranes where they were blocked in Odyssey Blocking Buffer with 0.1% (vol/vol) Tween 20. Rabbit anti-SipL_Δ LysM_ and mouse anti-SpoIVA (52) antibodies were used at a 1:2,500 dilution. Rabbit anti-mCherry (Abcam, Inc.) was used at a 1:2,000 dilution. IRDye 680CW and 800CW infrared dye-conjugated secondary antibodies were used at a 1:20,000 dilution, and blots were imaged on an Odyssey LiCor CLx imaging system.

### Spore purification

Sporulation was induced in 70:30 agar plates for 2 to 3 days as previously described (25, 53). *C. difficile* sporulating cells were washed 5-6 times in ice-cold water, incubated overnight at 4°C and treated with DNAse I (New England Biolabs) at 37°C for 60 minutes. Finally, spores were purified on a HistoDenz gradient (Sigma-Aldrich) and resuspended in water; spore purity was determined by phase-contrast microscopy (>95%), and the optical density of the spore preparation was measured at OD_600_. Spore yields were quantified by measuring the OD_600nm_ of the spore purifications from four 70:30 plates per replicate. The average of three biological replicates was calculated and statistical significance was determined using a one-way ANOVA and Tukey’s test. Spores were stored in water at 4°C.

### TEM analysis

Sporulating cultures (23-24 h) were fixed and processed for electron microscopy by the University of Vermont Microscopy Center as previously described (37). A minimum of 50 full-length sporulating cells were used for phenotype counting.

### mCherry fluorescence microscopy

Live-cell fluorescence microscopy was performed using Hoechst 33342 (15 μg/mL; Molecular Probes) and mCherry protein fusions. Samples were prepared on agarose pads either as previously described (21) or agarose pads prepared on Gene Frames (ThermoScientific). Briefly, one gene frame sticker was placed on a microscope slide, and 350 μL of 1% agarose was deposited inside the frame. Another microscope slide was set on top of the frame containing the agarose. The agarose pad was allowed to cool for 10 minutes at 4°C and then dried at room temperature for 5 minutes. Sporulating cultures were pipetted onto the agarose pad, covered with a cover-glass and imaged. Images were taken 30 minutes after harvesting the *C. difficile* sporulating cultures, since this allowed reconstitution of the mCherry fluorescence signal in an obligate anaerobe.

Phase-contrast and fluorescence microscopy were carried out on a Nikon 60x oil immersion objective (1.4 numerical aperture [NA]) using a Nikon 90i epifluorescence microscope. A CoolSnap HQ camera (Photometrics) was used to acquire multiple fields for each sample in 12-bit format with 2-by-2 binning, using NIS-Elements software (Nikon). The Texas Red channel was used to acquire images after 300 ms for mCherry-SpoIVA; 90 ms for SipL-mCherry; and 400 ms for CotE-mCherry. Hoechst stain was visualized using 90-ms exposure time and 3-ms exposure was used for phase-contrast pictures. Finally, 10 MHz images were imported to Adobe Photoshop CC 2017 software for pseudocoloring and minimal adjustments in brightness and contrast levels. Protein localization analysis were performed on a minimum of three independent biological replicates.

### Immunoprecipitation analyses

Immunoprecipitations with strains producing SipL-3xFLAG were performed on lysates prepared from cultures induced to sporulate on 70:30 plates for 24 hrs. The samples were processed as previously described (40), except that anti-FLAG magnetic beads (Sigma Aldrich) were used to pull-down FLAG-tagged proteins and any associated proteins. All immunoprecipitations were performed on three independent biological replicates.

### Bacterial two-hybrid analyses

Bacterial adenylate cyclase two-hybrid (BACTH) assays were performed using *E. coli* BTH101 cells based on the system first described by Karimova *et al*. (46). Briefly, for each assay, BTH101 cells were freshly co-transformed with ∼100 ng of each BACTH assay plasmid. The transformations were incubated on LB agar supplemented with 50 μg/mL kanamycin, 50 μg/mL carbenicillin, and 0.5 mM IPTG for 40 hrs at 30°C. After this incubation period, β-galactosidase activity was quantified in Miller units using the protocol described by (54, 55).

## Acknowledgments

We would like to thank N. Bishop and J. Schwarz for excellent assistance in preparing samples for transmission electron microscopy throughout this study; A. Camilli for access to the Nikon microscope; N. Minton (U. Nottingham) for generously providing us with access to the 630Δ*erm*Δ*pyrE* strain and pMTL-YN1C and pMTL-YN3 plasmids for allele-coupled exchange (ACE). Research in this manuscript was funded by R01AI22232 from the National Institutes of Allergy and Infectious Disease (NIAID) to A.S. A.S. is a Burroughs Wellcome Investigator in the Pathogenesis of Infectious Disease supported by the Burroughs Wellcome Fund. The content is solely the responsibility of the author(s) and does not necessarily reflect the views of the Burroughs Wellcome, NIAID, or the National Institutes of Health. The funders had no role in study design, data collection and interpretation, or the decision to submit the work for publication.

## Notes

### Competing Interest Statement

A. Shen is a paid consultant off BioVector, Inc.

## References

1. Abt MC, McKenney PT, Pamer EG. 2016. Clostridium difficile colitis: pathogenesis and host defence. Nat Rev Microbiol doi:10.1038/nrmicro.2016.108.

2. Smits WK, Lyras D, Lacy DB, Wilcox MH, Kuijper EJ. 2016. Clostridium difficile infection. Nat Rev Dis Primers 2:16020.

3. Chandrasekaran R, Lacy DB. 2017. The role of toxins in Clostridium difficile infection. FEMS Microbiol Rev 41:723–750.

4. Lewis BB, Pamer EG. 2017. Microbiota-Based Therapies for Clostridium difficile and Antibiotic-Resistant Enteric Infections. Annu Rev Microbiol 71:157–178.

5. Theriot CM, Young VB. 2015. Interactions Between the Gastrointestinal Microbiome and Clostridium difficile. Annu Rev Microbiol 69:445–61.

6. Paredes-Sabja D, Shen A, Sorg JA. 2014. Clostridium difficile spore biology: sporulation, germination, and spore structural proteins. Trends Microbiol 22:406–16.

7. Deakin LJ, Clare S, Fagan RP, Dawson LF, Pickard DJ, West MR, Wren BW, Fairweather NF, Dougan G, Lawley TD. 2012. The Clostridium difficile spooA gene is a persistence and transmission factor. Infect Immun 80:2704–11.

8. Dembek M, Willing SE, Hong HA, Hosseini S, Salgado PS, Cutting SM. 2017. Inducible Expression of spo0A as a Universal Tool for Studying Sporulation in Clostridium difficile. Front Microbiol 8:1793.

9. Swick MC, Koehler TM, Driks A. 2016. Surviving Between Hosts: Sporulation and Transmission. Microbiol Spectr 4.

10. Tan IS, Ramamurthi KS. 2014. Spore formation in Bacillus subtilis. Environ Microbiol Rep 6:212–25.

11. Shen A, Edwards AN, Sarker MR, Paredes-Sabja D. 2019. Sporulation and Germination in Clostridial Pathogens. Microbiol Spectr 7.

12. Popham DL, Bernhards CB. 2015. Spore Peptidoglycan. Microbiol Spectr 3.

13. Driks A, Eichenberger P. 2016. The Spore Coat. Microbiol Spectr 4.

14. Abecasis AB, Serrano M, Alves R, Quintais L, Pereira-Leal JB, Henriques AO. 2013. A genomic signature and the identification of new sporulation genes. J Bacteriol 195:2101–15.

15. Yutin N, Galperin MY. 2013. A genomic update on clostridial phylogeny: Gramnegative spore formers and other misplaced clostridia. Environ Microbiol 15:2631–2641.

16. Smith K, Bayer ME, Youngman P. 1993. Physical and functional characterization of the Bacillus subtilis spoIIM gene. J Bacteriol 175:3607–17.

17. Dembek M, Kelly A, Barwinska-Sendra A, Tarrant E, Stanley WA, Vollmer D, Biboy J, Gray J, Vollmer W, Salgado PS. 2018. Peptidoglycan degradation machinery in Clostridium difficile forespore engulfment. Mol Microbiol 110:390–410.

18. Ribis JW, Fimlaid KA, Shen A. 2018. Differential requirements for conserved peptidoglycan remodeling enzymes during Clostridioides difficile spore formation. Mol Microbiol 110:370–389.

19. Gutierrez J, Smith R, Pogliano K. 2010. SpoIID-mediated peptidoglycan degradation is required throughout engulfment during Bacillus subtilis sporulation. J Bacteriol 192:3174–86.

20. Doan T, Morlot C, Meisner J, Serrano M, Henriques A, Moran C, Rudner D. 2009. Novel secretion apparatus maintains spore integrity and developmental gene expression in Bacillus subtilis. PLoS Genet 5.

21. Fimlaid KA, Jensen O, Donnelly ML, Siegrist MS, Shen A. 2015. Regulation of Clostridium difficile Spore Formation by the SpoIIQ and SpoIIIA Proteins. PLoS Genet 11:e1005562.

22. Serrano M, Crawshaw AD, Dembek M, Monteiro JM, Pereira FC, Pinho MG, Fairweather NF, Salgado PS, Henriques AO. 2016. The SpoIIQ-SpoIIIAH complex of Clostridium difficile controls forespore engulfment and late stages of gene expression and spore morphogenesis. Mol Microbiol 100:204–28.

23. Londono-Vallejo JA, Frehel C, Stragier P. 1997. SpoIIQ, a forespore-expressed gene required for engulfment in Bacillus subtilis. Mol Microbiol 24:29–39.

24. Henriques AO, Moran CP, Jr. 2007. Structure, assembly, and function of the spore surface layers. Annu Rev Microbiol 61:555–88.

25. Ribis JW, Ravichandran P, Putnam EE, Pishdadian K, Shen A. 2017. The Conserved Spore Coat Protein SpoVM Is Largely Dispensable in Clostridium difficile Spore Formation. mSphere 2.

26. Levin PA, Fan N, Ricca E, Driks A, Losick R, Cutting S. 1993. An unusually small gene required for sporulation by Bacillus subtilis. Mol Microbiol 9:761–71.

27. Roels S, Driks A, Losick R. 1992. Characterization of spoIVA, a sporulation gene involved in coat morphogenesis in Bacillus subtilis. Journal of Bacteriology 174:575–585.

28. Ramamurthi KS, Lecuyer S, Stone HA, Losick R. 2009. Geometric cue for protein localization in a bacterium. Science 323:1354–7.

29. Tan IS, Weiss CA, Popham DL, Ramamurthi KS. 2015. A Quality-Control Mechanism Removes Unfit Cells from a Population of Sporulating Bacteria. Dev Cell 34:682–93.

30. McKenney PT, Eichenberger P. 2012. Dynamics of spore coat morphogenesis in Bacillus subtilis. Mol Microbiol 83:245–60.

31. Gill RL, Jr., Castaing JP, Hsin J, Tan IS, Wang X, Huang KC, Tian F, Ramamurthi KS. 2015. Structural basis for the geometry-driven localization of a small protein. Proc Natl Acad Sci U S A 112:E1908–15.

32. Ramamurthi KS, Clapham KR, Losick R. 2006. Peptide anchoring spore coat assembly to the outer forespore membrane in Bacillus subtilis. Mol Microbiol 62:1547–1557.

33. Peluso EA, Updegrove TB, Chen J, Shroff H, Ramamurthi KS. 2019. A 2-dimensional ratchet model describes assembly initiation of a specialized bacterial cell surface. Proc Natl Acad Sci U S A 116:21789–21799.

34. Castaing J-P, Nagy A, Anantharaman V, Aravind L, Ramamurthi K. 2013. ATP hydrolysis by a domain related to translation factor GTPases drives polymerization of a static bacterial morphogenetic protein. Proc Natl Acad Sci U S A 110:60.

35. Ramamurthi KS, Losick R. 2008. ATP-driven self-assembly of a morphogenetic protein in Bacillus subtilis. Mol Cell 31:406–14.

36. Wang KH, Isidro AL, Domingues L, Eskandarian HA, McKenney PT, Drew K, Grabowski P, Chua MH, Barry SN, Guan M, Bonneau R, Henriques AO, Eichenberger P. 2009. The coat morphogenetic protein SpoVID is necessary for spore encasement in Bacillus subtilis. Mol Microbiol 74:634–49.

37. Putnam EE, Nock AM, Lawley TD, Shen A. 2013. SpoIVA and SipL are Clostridium difficile spore morphogenetic proteins. J Bacteriol 195:1214–25.

38. Decker AR, Ramamurthi KS. 2017. Cell Death Pathway That Monitors Spore Morphogenesis. Trends Microbiol doi:10.1016/j.tim.2017.03.005.

39. Ebmeier SE, Tan IS, Clapham KR, Ramamurthi KS. 2012. Small proteins link coat and cortex assembly during sporulation in Bacillus subtilis. Mol Microbiol 84:682–96.

40. Touchette MH, Benito de la Puebla H, Ravichandran P, Shen A. 2019. SpoIVA-SipL Complex Formation Is Essential for Clostridioides difficile Spore Assembly. J Bacteriol 201.

41. Castaing JP, Lee S, Anantharaman V, Ravilious GE, Aravind L, Ramamurthi KS. 2014. An autoinhibitory conformation of the Bacillus subtilis spore coat protein SpoIVA prevents its premature ATP-independent aggregation. FEMS Microbiol Lett 358:145–53.

42. Bennison DJ, Irving SE, Corrigan RM. 2019. The Impact of the Stringent Response on TRAFAC GTPases and Prokaryotic Ribosome Assembly. Cells 8.

43. Ng YK, Ehsaan M, Philip S, Collery MM, Janoir C, Collignon A, Cartman ST, Minton NP. 2013. Expanding the repertoire of gene tools for precise manipulation of the Clostridium difficile genome: allelic exchange using pyrE alleles. PLoS One 8:e56051.

44. Shen A, Fimlaid KA, Pishdadian K. 2016. Inducing and Quantifying Clostridium difficile Spore Formation. Methods Mol Biol 1476:129–42.

45. Hong HA, Ferreira WT, Hosseini S, Anwar S, Hitri K, Wilkinson AJ, Vahjen W, Zentek J, Soloviev M, Cutting SM. 2017. The Spore Coat Protein CotE Facilitates Host Colonisation by Clostridium difficile. J Infect Dis doi:10.1093/infdis/jix488.

46. Karimova G, Pidoux J, Ullmann A, Ladant D. 1998. A bacterial two-hybrid system based on a reconstituted signal transduction pathway. Proc Natl Acad Sci U S A 95:5752–6.

47. Beall B, Driks A, Losick R, Moran CP, Jr. 1993. Cloning and characterization of a gene required for assembly of the Bacillus subtilis spore coat. J Bacteriol 175:1705–16.

48. Srikhanta YN, Hutton ML, Awad MM, Drinkwater N, Singleton J, Day SL, Cunningham BA, McGowan S, Lyras D. 2019. Cephamycins inhibit pathogen sporulation and effectively treat recurrent Clostridioides difficile infection. Nat Microbiol 4:2237–2245.

49. Sorg JA, Dineen SS. 2009. Laboratory maintenance of Clostridium difficile. Curr Protoc Microbiol Chapter 9:Unit 9A 1.

50. Karasawa T, Ikoma S, Yamakawa K, Nakamura S. 1995. A defined growth medium for Clostridium difficile. Microbiology 141 (Pt 2):371–5.

51. Donnelly ML, Li W, Li YQ, Hinkel L, Setlow P, Shen A. 2017. A Clostridium difficile-Specific, Gel-Forming Protein Required for Optimal Spore Germination. mBio 8.

52. Kevorkian Y, Shirley DJ, Shen A. 2015. Regulation of Clostridium difficile spore germination by the CspA pseudoprotease domain. Biochimie doi:10.1016/j.biochi.2015.07.023.

53. Diaz OR, Sayer CV, Popham DL, Shen A. 2018. Clostridium difficile Lipoprotein GerS Is Required for Cortex Modification and Thus Spore Germination. mSphere 3.

54. Dahlstrom KM, Giglio KM, Collins AJ, Sondermann H, O’Toole GA. 2015. Contribution of Physical Interactions to Signaling Specificity between a Diguanylate Cyclase and Its Effector. mBio 6:e01978–15.

55. Giacalone D, Smith TJ, Collins AJ, Sondermann H, Koziol LJ, O’Toole GA. 2018. Ligand-Mediated Biofilm Formation via Enhanced Physical Interaction between a Diguanylate Cyclase and Its Receptor. mBio 9.

56. Fimlaid KA, Bond JP, Schutz KC, Putnam EE, Leung JM, Lawley TD, Shen A. 2013. Global Analysis of the Sporulation Pathway of Clostridium difficile. PLoS Genet 9:e1003660.

57. Degasperi A, Birtwistle MR, Volinsky N, Rauch J, Kolch W, Kholodenko BN. 2014. Evaluating strategies to normalise biological replicates of Western blot data. PLoS One 9:e87293.

