## Supplementary Materials for "Role of SpoIVA ATPase Motifs During *Clostridioides difficile* Sporulation"

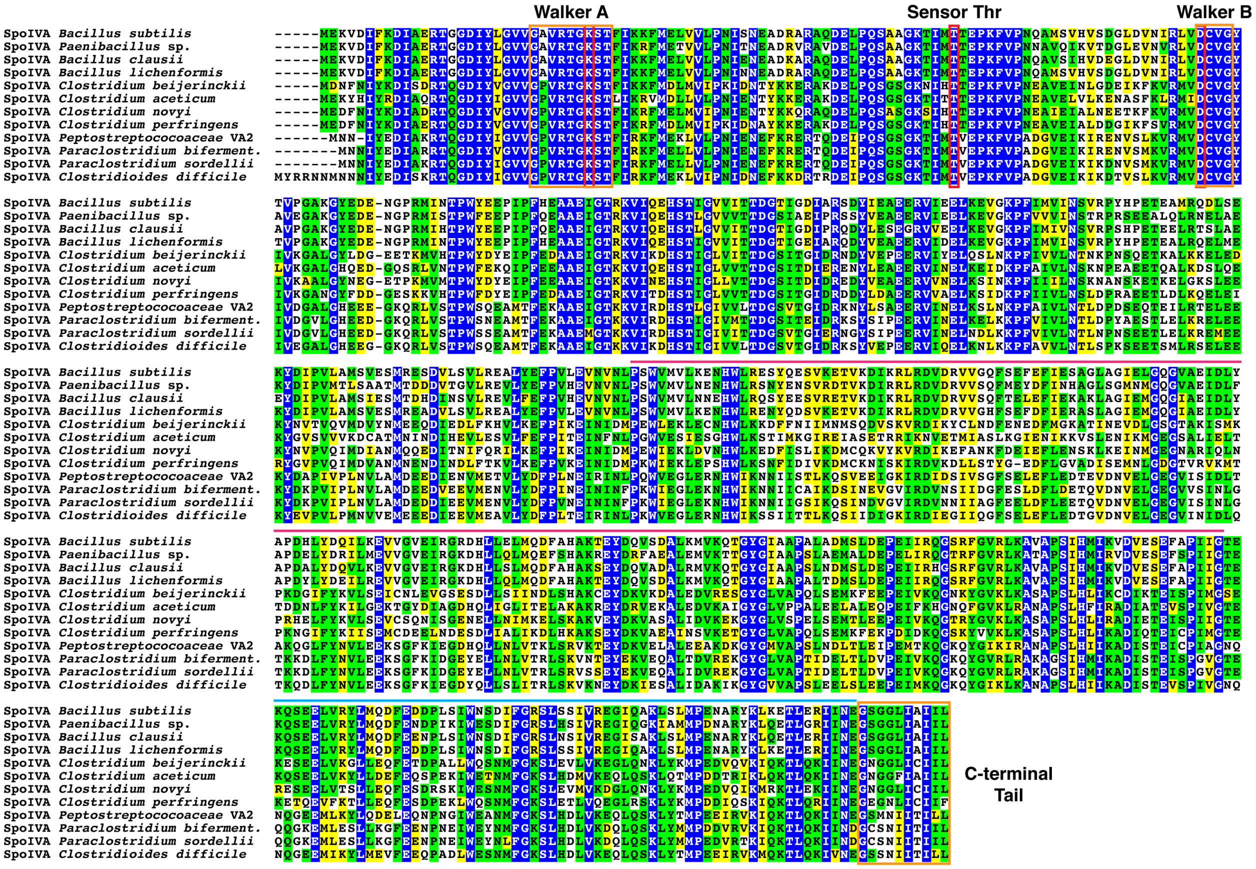


**Figure S1. Sequence alignment of SpoIVA homologs highlighting ATPase motifs**. The Walker A motif is required for ATP binding, while the Sensor Threonine and Walker B motifs are required for ATP hydrolysis (1). The motifs are boxed in orange as is the C-terminal tail region, which has been implicated in binding to SpoVM (2). The pink line highlights the central domain, which is required for the conformational changes induced by ATP hydrolysis in *B. subtilis* SpoIVA (3). The blue line highlights the C-terminal domain globular domain defined by Castaing *et al*. (4). The accession numbers for the SpoIVA homologs are given in parentheses: *B. subtilis* 168 (NP_390161), *Paenibacillus* sp. GYMC10_2192 (YP_003242280), *B. clausii* YP_175378, *B. lichenformis* (YP_079580), *Clostridium beijerinckii* (WP_02688916), *C. aceticum* (WP_044825087), *C. novyi* (YP_878326), *C. perfringens* (NP_562669), *Peptostreptococcaceae* VA2 (WP_026900372), *Paraclostridium bifermentans*

(WP_021433867), *Paraclostridium sordellii* (CEQ01088), and *Clostridioides difficile* (YP_001089140).

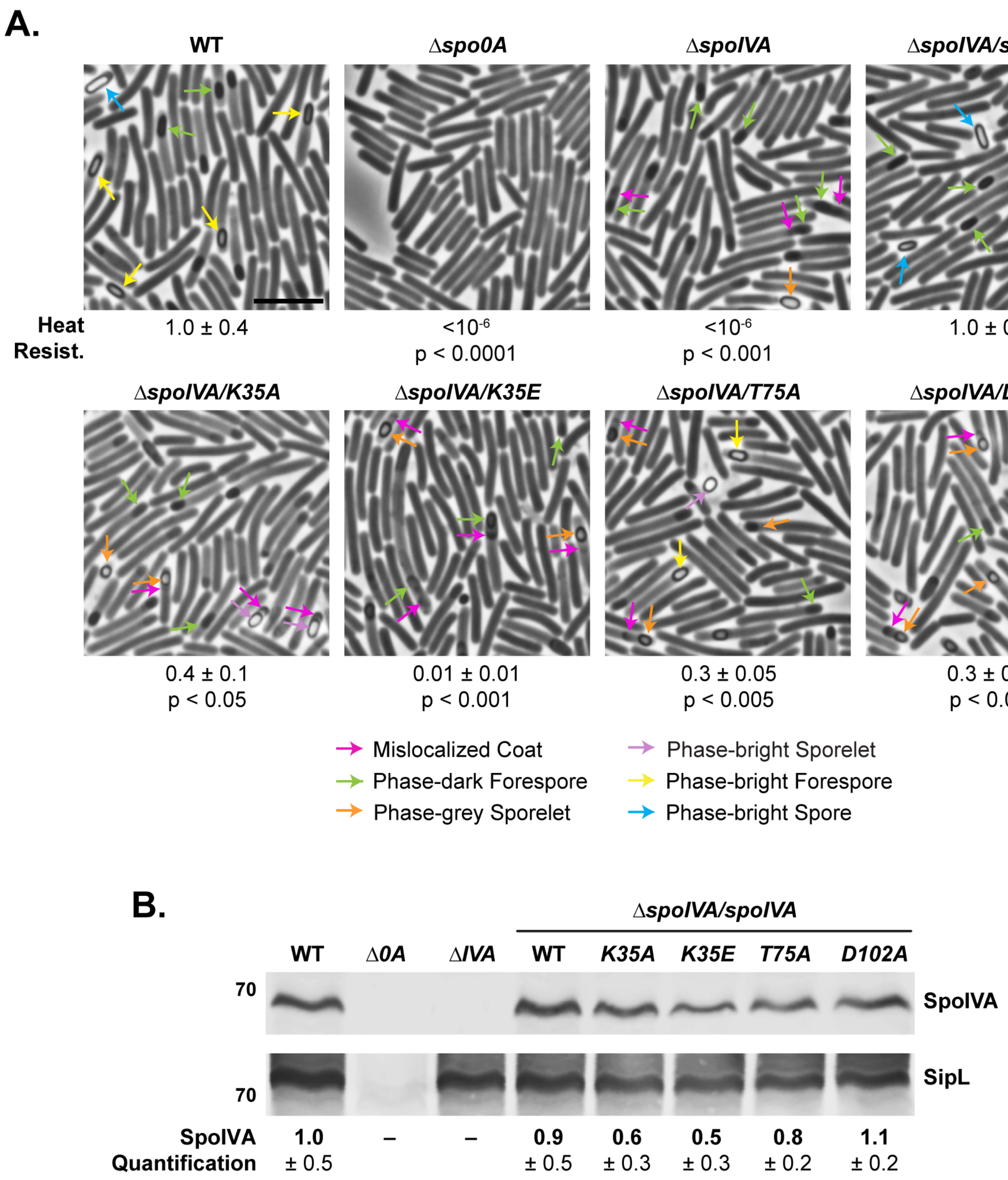
A.
WT
Δspo0A
ΔspoIVA
ΔspoIVA/spoIVA
Heat Resist.
1.0 ± 0.4
<10^-6
p < 0.0001
<10^-6
p < 0.001
1.0 ± 0.7
ΔspoIVA/K35A
ΔspoIVA/K35E
ΔspoIVA/T75A
ΔspoIVA/D102A
0.4 ± 0.1
p < 0.05
0.01 ± 0.01
p < 0.001
0.3 ± 0.05
p < 0.005
0.3 ± 0.1
p < 0.01
Mislocalized Coat
Phase-dark Forespore
Phase-grey Sporelet
Phase-bright Sporelet
Phase-bright Forespore
Phase-bright Spore
B.
ΔspoIVA/spoIVA
WT
Δ0A
ΔIVA
WT
K35A
K35E
T75A
D102A
70
SpoIVA
SipL
70
SpoIVA Quantification
1.0 ± 0.5
–
–
0.9 ± 0.5
0.6 ± 0.3
0.5 ± 0.3
0.8 ± 0.2
1.1 ± 0.2

**Figure S2. Effect of SpoIVA ATPase motif mutations encoded in the ectopic *pyrE* locus on functional spore formation**. (A) Phase-contrast microscopy analyses of the indicated *C. difficile* strains ~20 hrs after sporulation induction. Arrows mark examples of sporulating cells at different stages of maturation: pink arrows mark regions of mislocalized coat based on previous studies (5, 6); green arrows highlight immature phase-dark forespores; orange arrows highlight phase-gray sporelets, which look swollen and are surrounded by a phase-dark ring; purple arrows highlight phase-bright sporelets, which are swollen and surrounded by a phase-dark ring; yellow arrows mark mature phase-bright forespores (phase-brightness reflects cortex formation (7, 8)); blue arrows highlight phase-bright free spores. Heat resistance efficiencies are based on 20-24 hr sporulating cultures and represent the mean and standard deviation for a given strain relative to wild type based on a minimum of three biological replicates. Statistical significance for all assays was determined relative to wild type using a one-way ANOVA and Tukey's test. Scale bar represents 5 μm. The limit of detection of the assay is $10^{-6}$. (B) Western blot analyses of SpoIVA and SipL. SpoIVA levels were quantified based on analyses of three biological replicates using (9). Statistical significance for all assays was determined relative to wild type using a one-way ANOVA and Tukey's test. No statistically significant differences were detected by western blotting using a one-way ANOVA and Tukey's test.

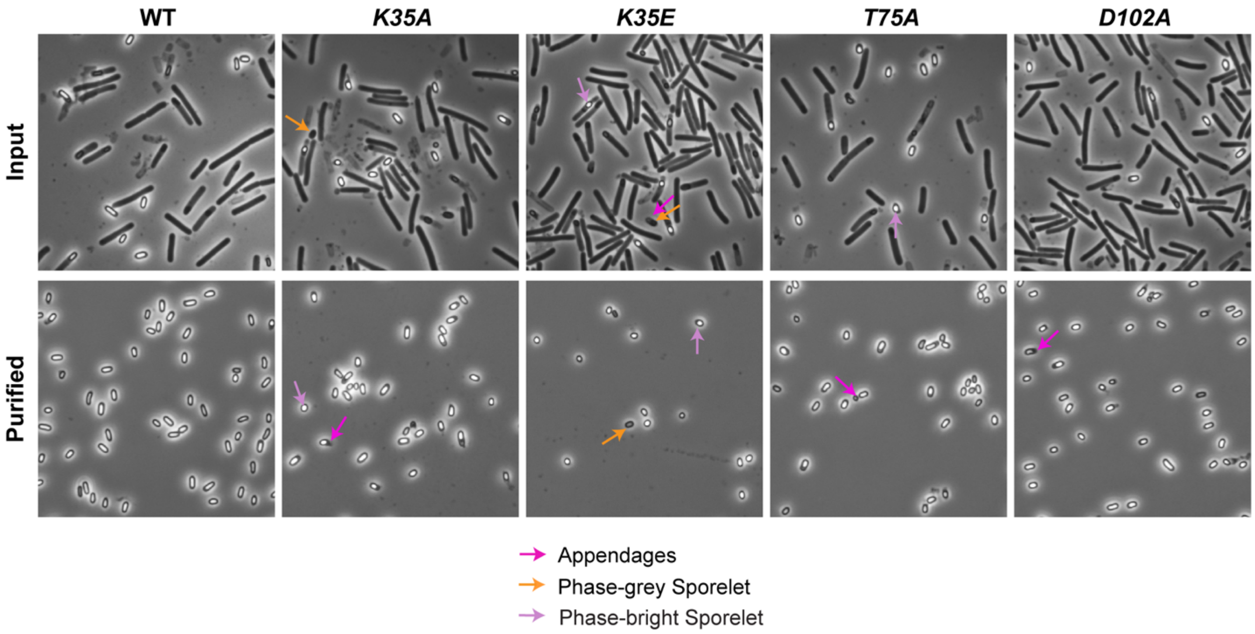


**Figure S3. Spore purifications of SpoIVA ATPase motif mutants**. The "Input" sample was taken after cells were scraped off sporulation media and resuspended in ice-cold water. The "Purified" sample was taken after spores were washed repeatedly and purified on a density gradient. Pink arrows mark regions of probable coat attachments or "appendages" (https://www.biorxiv.org/content/10.1101/468637v1); orange arrows highlight phase-gray sporelets, which look swollen and are surrounded by a phase-dark ring; purple arrows highlight phase-bright sporelets, which are swollen and surrounded by a phase-dark ring.

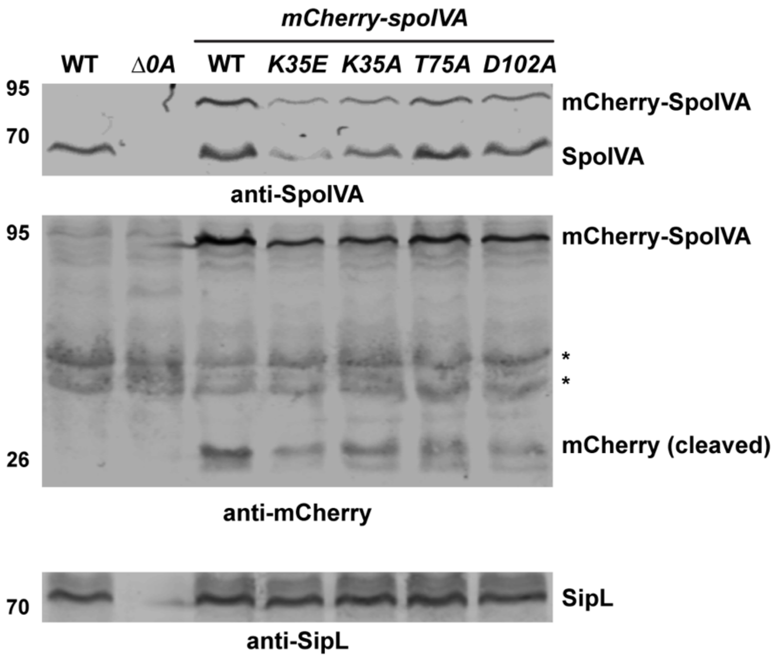


**Figure S4. Western blot analyses of mCherry-SpoIVA levels in SpoIVA ATPase motif mutants**. Antibodies to SpoIVA, mCherry, and SipL were used as indicated. Asterisks indicate non-specific bands bound by the mCherry antibody. The western blots are representative of the results of two biological replicates.

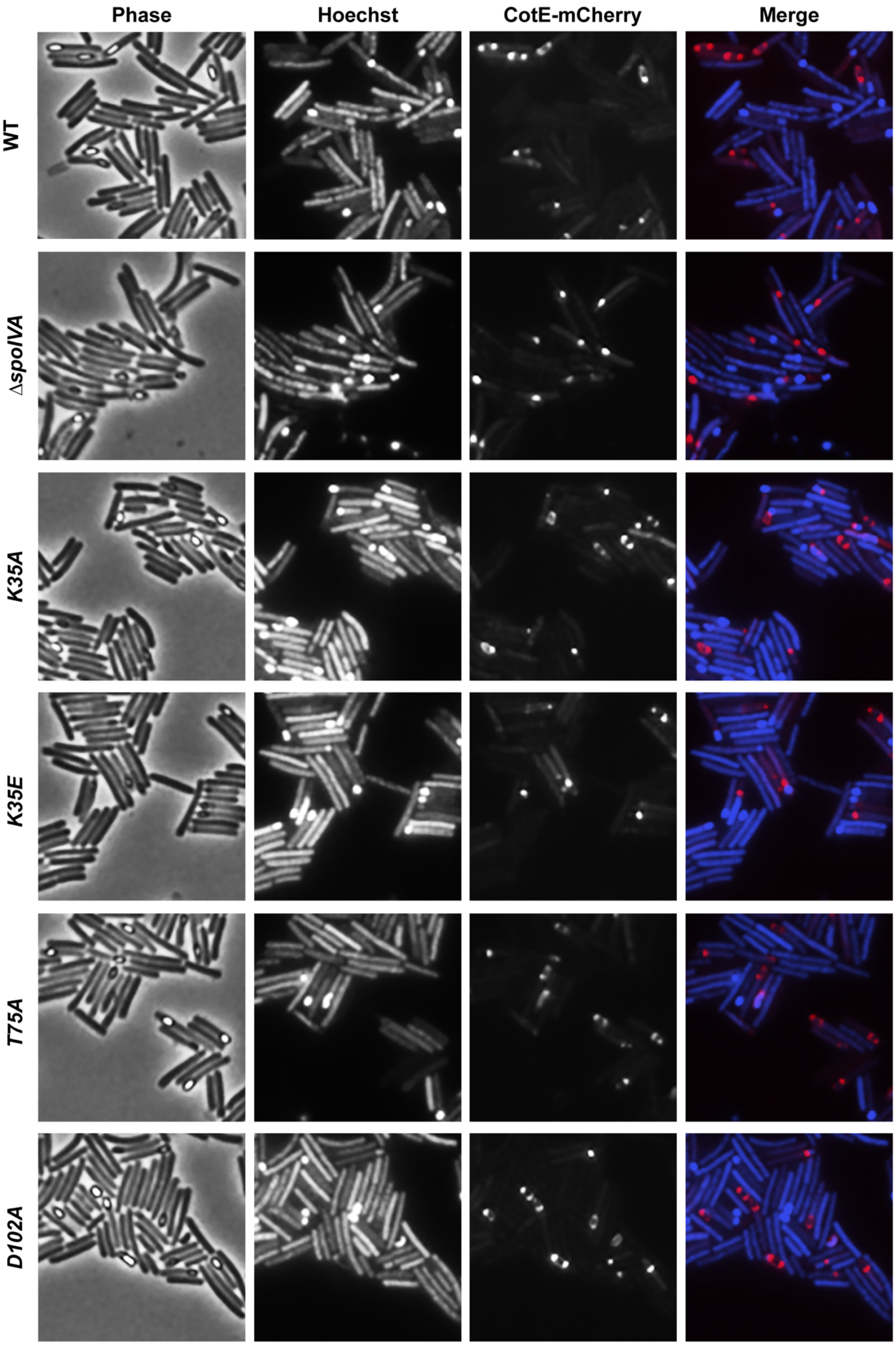

Phase
Hoechst
CotE-mCherry
Merge
WT
ΔspoIVA
K35A
K35E
T75A
D102A

**Figure S5. Effect of SpoIVA ATPase motif mutations on CotE localization**. Fluorescence microscopy analyses of WT and *spoIVA* mutants encoding ATPase motif mutations expressing *cotE-mCherry* from the *pyrE* locus. Microscopy was performed on samples 23 hrs after sporulation induction. Phase-contrast microscopy was used to visualize sporulating cells (Phase). Hoechst staining used to visualize the nucleoid is shown in blue, and CotE-mCherry fluorescence is shown in red. The merge of the Hoechst staining and mCherry signal is shown. Images are representative of the results of three biological replicates.

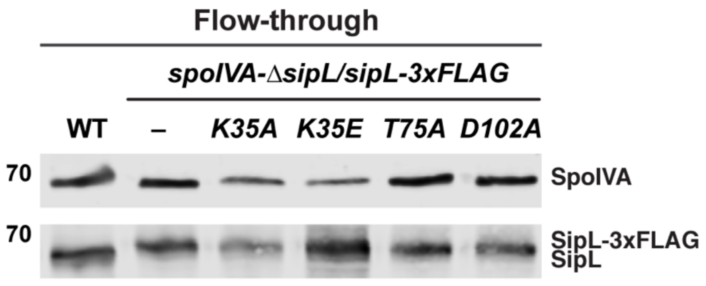


**Figure S6. Western blot analysis of the flow-through fraction of co-immunoprecipitations of SipL-3xFLAG in SpoIVA ATPase motif mutants.** SipL-3xFLAG was immunoprecipitated from cleared lysates prepared from either wild type (WT), Δ*sipL/sipL-3xFLAG* complementation strain (–), or Δ*sipL/sipL-3xFLAG* strains encoding SpoIVA ATPase motif mutations in their native locus. The "Flow-through" sample was taken from the supernatant after anti-FLAG magnetic beads were pelleted. It represents proteins that were either unable to bind to the beads specifically or were present in excess of the binding capacity of the beads. The SpoIVA present in the WT flow-through sample analysis indicates that the anti-FLAG beads were saturated.

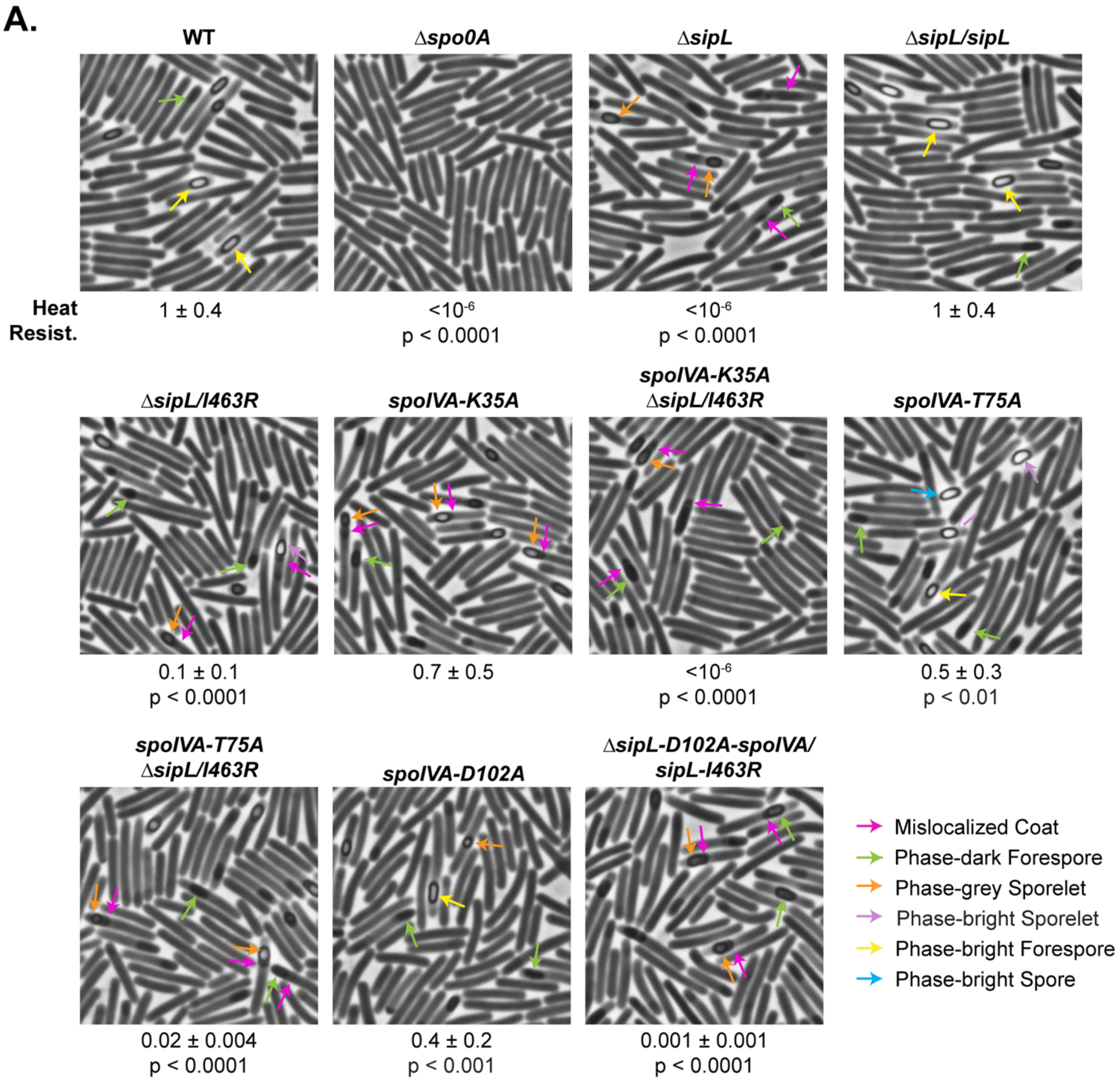
A.
WT
Δspo0A
ΔsipL
ΔsipL/sipL
Heat
Resist.
1 ± 0.4
<10^-6
p < 0.0001
<10^-6
p < 0.0001
1 ± 0.4
ΔsipL/I463R
spoIVA-K35A
spoIVA-K35A
ΔsipL/I463R
spoIVA-T75A
0.1 ± 0.1
p < 0.0001
0.7 ± 0.5
<10^-6
p < 0.0001
0.5 ± 0.3
p < 0.01
spoIVA-T75A
ΔsipL/I463R
spoIVA-D102A
ΔsipL-D102A-spoIVA/
sipL-I463R
0.02 ± 0.004
p < 0.0001
0.4 ± 0.2
p < 0.001
0.001 ± 0.001
p < 0.0001
Mislocalized Coat
Phase-dark Forespore
Phase-grey Sporelet
Phase-bright Sporelet
Phase-bright Forespore
Phase-bright Spore

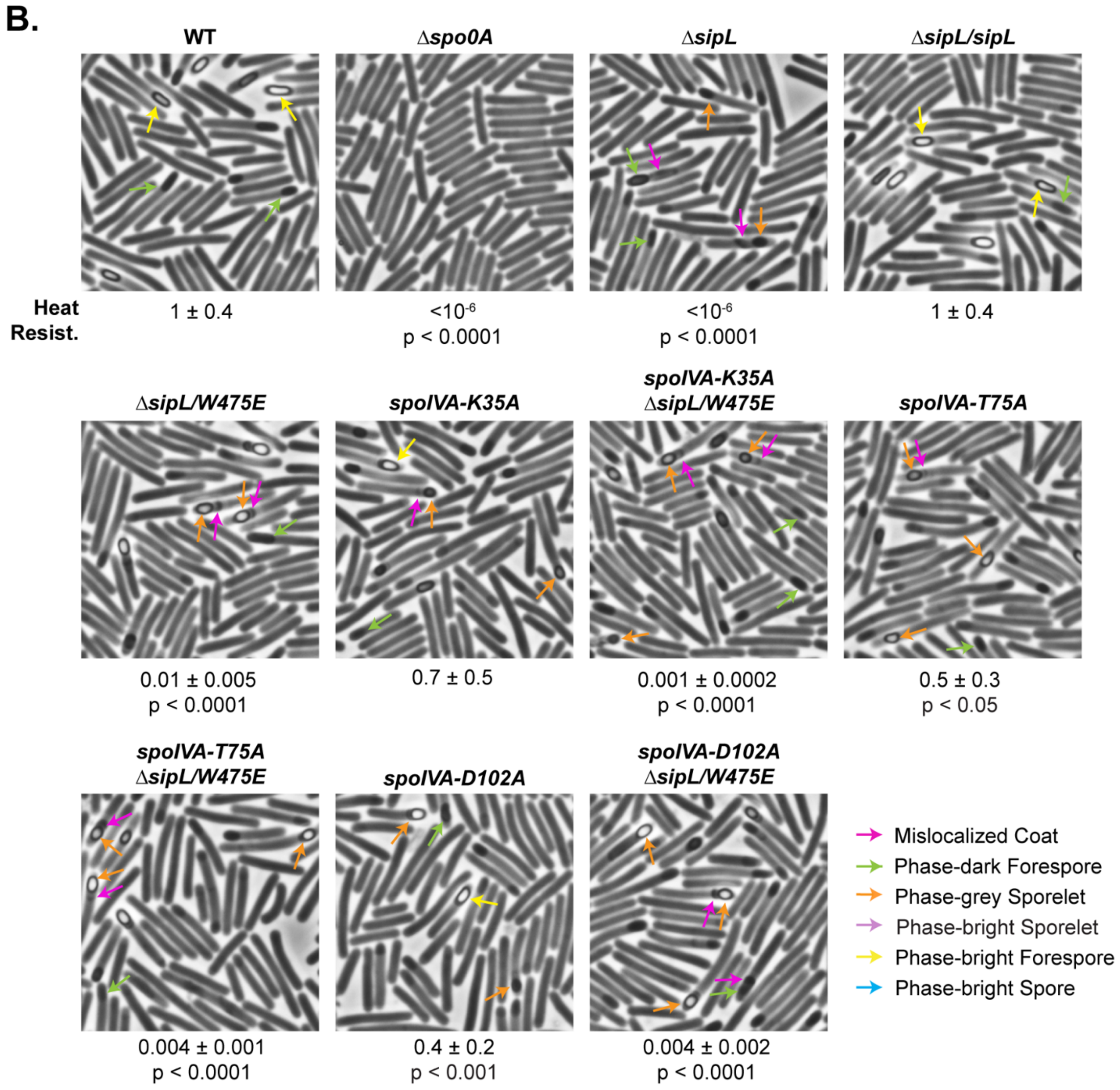


**Figure S7. Effect of combining SpoIVA ATPase motif mutations with *sipL* (A) *I463R* and (B) *W475E* mutations**. Phase-contrast microscopy analyses of the indicated *C. difficile* strains ~20 hrs after sporulation induction. The SpoIVA ATPase motif mutations are encoded in the native *spoIVA* locus, while the SipL LysM domain mutations are encoded in a *sipL* gene integrated into the *pyrE* locus of a Δ*sipL* strain. Arrows mark examples of sporulating cells at different stages of maturation: pink arrows mark regions of mislocalized coat based on previous studies (5, 6); green arrows highlight immature phase-dark forespores; orange arrows highlight phase-gray sporelets, which look swollen and are surrounded by a phase-dark ring; purple arrows highlight phase-bright sporelets, which are swollen and surrounded by a phase-dark ring; yellow arrows mark mature phase-bright forespores (phase-brightness reflects cortex formation (7, 8)); blue arrows highlight phase-bright free spores. Heat resistance efficiencies are based on 20-24 hr sporulating cultures and represent the mean and standard deviation for a given strain relative to wild type based on a minimum of three biological replicates. The limit of detection of the assay is $10^{-6}$. Statistical significance was determined relative to wild type using a one-way ANOVA and Tukey's test.

**Supplementary Text S1 – *E. coli* strain construction**

**pMTL-YN1C-*spoIVA* ATPase motif mutations**. To clone the *K35A spoIVA pyrE* locus complementation construct, primer pair #2036 and 1483 was used to amplify the region 117 bp upstream of *spoIVA* up to the region around the K35 codon off *C. difficile* genomic DNA. Primer pair #1482 and 2037 was used to amplify the *spoIVA* gene starting from the region around the K35 codon through to the stop codon off *C. difficile* genomic DNA. Primers 1482 and 1483 encode the K35A mutation. The resulting PCR products were cloned into pMTL-YN1C digested with NotI and XhoI using Gibson assembly, although a SOE PCR step was sometimes employed to join the two *spoIVA* fragments together (10). The same procedure was used to clone the *K35E*, *T75A*, and *D102A* mutations except primer pairs #711 and 712; #1215 and 1216; and #1217 and 1218, respectively were used to introduce the indicated mutations.

**pMTL-YN3-*spoIVA* ATPase motif mutations.** Primer pair #1927 and 1483 were used to amplify the region 1043 bp upstream of the *spoIVA* gene through to the K35 codon off *C. difficile* genomic DNA. Primer pair #1482 and 1928 were used to amplify the region downstream of the K35 codon through to 975 bp downstream of the *spoIVA* gene off *C. difficile* genomic DNA, although a SOE PCR step was sometimes employed to join the two *spoIVA* fragments together (10). The PCR products resulting PCR products were cloned into pMTL-YN3 digested with AscI and SbfI using Gibson assembly.

**pMTL-YN1C-*mCherry-spoIVA* ATPase motif mutations**. To generate a construct encoding mCherry fusions to SpoIVA ATPase motif mutants, primer pair #2036 and 1483 was used to amplify the promoter region of *spoIVA*, a codon-optimized *mCherry* gene up to the region around the K35 codon using pMTL-YN1C *mCherry-spoIVA* as the template (6). Primer pair #1482 and 2037 was used to amplify the *spoIVA* gene starting from the region around the K35 codon through to the stop codon off *C. difficile* genomic DNA. The resulting PCR products were gel purified and used in a PCR SOE reaction (10) to generate the *mCherry-spoIVA* ATPase motif mutant constructs, which were assembled into pMTL-YN1C using Gibson assembly. The same procedure was used to clone the *K35E*, *T75A*, and *D102A* mutations except primer pairs #711 and 712; #1215 and 1216; and #1217 and 1218, respectively were used to introduce the indicated mutations.

**pKNT25-*sipL***. Primer pair #1453 and 1454 were used to amplify the *sipL* gene (without the stop codon) with BamHI and KpnI sites flanking the 5' and 3' ends of the gene. The resulting PCR product was digested with BamHI and KpnI then ligated into pKNT25 digested with the same enzymes (11). The ligation was transformed into DH5α. The resulting plasmid encodes *sipL* as an N-terminal fusion to the T25 fragment of adenylate cyclase.

**pUT18C-*spoIVA***. Primer pair #1452 and 1337 were used to amplify the *spoIVA* gene including the stop codon with BamHI and KpnI sites flanking the 5' and 3' ends of the gene. The resulting PCR product was digested with BamHI and KpnI then ligated into pUT18C digested with the same enzymes (11). The ligation was transformed into DH5α. The resulting plasmid encodes *spoIVA* as a C-terminal fusion to the T18 fragment of adenylate cyclase.

**pUT18C-*spoIVA* ATPase motif mutations**. Primer pair #3064 and #3065 was used to amplify *spoIVA* encoding ATPase motif mutations off pMTL-YN1C *spoIVA K35A*, *K35E*, *T75A*, and *D102A*, respectively. The resulting PCR products were cloned into pUT18C digested with BamHI and KpnI.

### Supplementary Table S1. Strains used in this study

**_C. difficile_ strains – 630Δ_erm_**

| Strain # | Strain name | Relevant genotype or features | Source/reference |
|---|---|---|---|
| 803 | 630Δ*erm*Δ*pyrE* Δ*spoIVA* | 630Δ*erm* Δ*pyrE* with *spoIVA* (*CD2629*) deleted | (6) |
| 846 | 630Δ*erm*-p | *erm*-sensitive derivate of 630 with *pyrE* restored | (12) |
| 849 | 630Δ*erm* Δ*spo0A*-p | 630Δ*erm* Δ*spo0A* with *pyrE* restored | (12) |
| 880 | 630Δ*erm* Δ*spoIVA*-p | 630Δ*erm* Δ*spoIVA* with *pyrE* restored | (6) |
| 883 | 630Δ*erm* Δ*spoIVA*/ *spoIVA* | 630Δ*erm* Δ*spoIVA* with *spoIVA* in the *pyrE* locus | (6) |
| 886 | 630Δ*erm* Δ*spoIVA*/ *spoIVA*$_{K35A}$ | 630Δ*erm* Δ*spoIVA* with *spoIVA*$_{K35A}$ in the *pyrE* locus | This study |
| 892 | 630Δ*erm* Δ*spoIVA*/ *spoIVA*$_{T75A}$ | 630Δ*erm* Δ*spoIVA* with *spoIVA*$_{T75A}$ in the *pyrE* locus | This study |
| 896 | 630Δ*erm* Δ*spoIVA*/ *spoIVA*$_{D102A}$ | 630Δ*erm* Δ*spoIVA* with *spoIVA*$_{D102A}$ in the *pyrE* locus | This study |
| 968 | 630Δ*erm* Δ*spoIVA*/ *spoIVA*$_{K35E}$ | 630Δ*erm*Δ*spoIVA* with *spoIVA*$_{K35E}$ in the *pyrE* locus | This study |
| 1010 | 630Δ*erm* Δ*sipL*-p | 630Δ*erm* with *sipL* deleted and *pyrE* restored | (13) |
| 1013 | 630Δ*erm* Δ*sipL*/*sipL* | 630Δ*erm* Δ*sipL* with *sipL* in the *pyrE* locus | (13) |
| 1144 | 630Δ*erm*/*mCherry-IVA* | 630Δ*erm* with *mCherry-IVA* in the *pyrE* locus | (6) |
| 1158 | 630Δ*erm* Δ*sipL*/*sipL-mCherry* | 630Δ*erm* Δ*sipL* with *sipL-mCherry* in the *pyrE* locus | (13) |
| 1289 | 630Δ*erm* Δ*pyrE spoIVA-D102A* | 630Δ*erm* Δ*pyrE* Δ*spoIVA* with *D102A* in the *spoIVA* locus | This study |
| 1295 | 630Δ*erm spoIVA-D102A*/ *mCherry-spoIVA*$_{D102A}$ | 630Δ*erm* Δ*spoIVA* with *D102A* in the *spoIVA* locus and *mCherry-spoIVA*$_{D102A}$ in the *pyrE* locus | This study |
| 1306 | 630Δ*erm*/ *cotE-mCherry* | 630Δ*erm* with *cotE-mCherry* in the *pyrE* locus | (13) |
| 1331 | 630Δ*erm spoIVA-D102A*-p | 630Δ*erm* Δ*spoIVA* with *D102A* in the *spoIVA* locus and *pyrE* restored | This study |
| 1334 | 630Δ*erm* Δ*pyrE spoIVA-K35A* | 630Δ*erm* Δ*pyrE* with *K35A* in the *spoIVA* locus | This study |
| 1337 | 630Δ*erm* Δ*pyrE spoIVA-K35E* | 630Δ*erm* Δ*pyrE* with *K35E* in the *spoIVA* locus | This study |
| 1340 | 630Δ*erm* Δ*pyrE spoIVA-T75A* | 630Δ*erm* Δ*pyrE* with *T75A* in the *spoIVA* locus | This study |
| 1343 | 630Δ*erm spoIVA-K35E*-p | 630Δ*erm* with *K35E* in the *spoIVA* locus and *pyrE* restored | This study |
| 1346 | 630Δ*erm spoIVA-K35E*/ *mCherry-spoIVA*$_{K35E}$ | 630Δ*erm* with *K35E* in the *spoIVA* locus and *mCherry-spoIVA*$_{K35E}$ in the *pyrE* locus | This study |
| 1354 | 630Δ*erm spoIVA-K35A*-p | 630Δ*erm* with *K35A* in the *spoIVA* locus and *pyrE* restored | This study |
| 1357 | 630Δ*erm spoIVA-K35A*/ *mCherry-spoIVA*$_{K35A}$ | 630Δ*erm* with *K35A* in the *spoIVA* locus and *mCherry-spoIVA*$_{K35A}$ in the *pyrE* locus | This study |
| 1377 | 630Δ*erm* Δ*sipL*/*sipL-3XFLAG* | 630Δ*erm* Δ*sipL* with *sipL-3XFLAG* in the *pyrE* locus | (14) |
| 1396 | 630Δ*erm spoIVA-T75A*-p | 630Δ*erm* with *T75A* in the *spoIVA* locus and *pyrE* restored | This study |
| 1399 | 630Δ*erm spoIVA-T75A*/ *mCherry-spoIVA*$_{T75A}$ | 630Δ*erm* with *T75A* in the *spoIVA* locus and *mCherry-spoIVA*$_{T75A}$ in the *pyrE* locus | This study |
| 1456 | 630Δ*erm* Δ*sipL*/*sipL*$_{I463R}$ | 630Δ*erm*Δ*sipL* with *sipL*$_{I463R}$ in the *pyrE* locus | (14) |
| 1500 | 630Δ*erm* Δ*sipL*/*sipL*$_{W475E}$ | 630Δ*erm*Δ*sipL* with *sipL*$_{W475E}$ in the *pyrE* locus | (14) |
| 1852 | 630Δ*erm* Δ*sipL*/*sipL-3XFLAG* | 630Δ*erm* Δ*sipL* with *sipL-3XFLAG* in the *pyrE* locus | (14) |
| 2436 | 630Δ*erm* Δ*pyrE* Δ*sipL spoIVA-K35A* | 630Δ*erm* Δ*pyrE* Δ*sipL* with *K35A* in the *spoIVA* locus | This study |
| 2440 | 630Δ*erm* Δ*pyrE* Δ*sipL spoIVA-K35E* | 630Δ*erm* Δ*pyrE* Δ*sipL* Δ*pyrE* with *K35E* in the *spoIVA* locus | This study |
| 2457 | 630Δ*erm* Δ*pyrE* Δ*sipL spoIVA-T75A* | 630Δ*erm* Δ*pyrE* Δ*sipL* with *T75A* in the *spoIVA* locus | This study |
| 2460 | 630Δ*erm* Δ*pyrE* Δ*sipL spoIVA-D102A* | 630Δ*erm* Δ*pyrE* Δ*sipL* with *D102A* in the *spoIVA* locus | This study |
| 2466 | 630Δ*erm* Δ*sipL spoIVA-K35A*/ *sipL-3xFLAG* | 630Δ*erm* Δ*sipL* with *K35A* in the *spoIVA* locus and *sipL-3xFLAG* in the *pyrE* locus | This study |
| 2469 | 630Δ*erm* Δ*sipL spoIVA-K35E*/ *sipL-3xFLAG* | 630Δ*erm* Δ*sipL* with *K35E* in the *spoIVA* locus and *sipL-3xFLAG* in the *pyrE* locus | This study |
| 2472 | 630Δ*erm* Δ*sipL spoIVA-T75A*/ *sipL-3xFLAG* | 630Δ*erm* Δ*sipL* with *T75A* in the *spoIVA* locus and *sipL-3xFLAG* in the *pyrE* locus | This study |

| | | | |
|---|---|---|---|
| 2475 | 630Δ*erm* Δ*sipL spoIVA-D102A/ sipL-3xFLAG* | 630Δ*erm* Δ*sipL* with *D102A* in the *spoIVA* locus and *sipL-3xFLAG* in the *pyrE* locus | This study |
| 2532 | 630Δ*erm* Δ*spoIVA /cotE-mCherry* | 630Δ*erm* Δ*spoIVA* with *cotE-mCherry* in the *pyrE* locus | This study |
| 2538 | 630Δ*erm spoIVA-K35E/ cotE-mCherry* | 630Δ*erm* with *K35E* in the *spoIVA* locus and *cotE-mCherry* in the *pyrE* locus | This study |
| 2545 | 630Δ*erm spoIVA-K35A/ cotE-mCherry* | 630Δ*erm* with *K35A* in the *spoIVA* locus and *cotE-mCherry* in the *pyrE* locus | This study |
| 2548 | 630Δ*erm spoIVA-T75A/ cotE-mCherry* | 630Δ*erm* with *T75A* in the *spoIVA* locus and *cotE-mCherry* in the *pyrE* locus | This study |
| 2642 | 630Δ*erm spoIVA-D102A/ cotE-mCherry* | 630Δ*erm* with *D102A* in the *spoIVA* locus and *cotE-mCherry* in the *pyrE* locus | This study |
| 2659 | 630Δ*erm* Δ*sipL spoIVA-K35A/ sipL-mCherry* | 630Δ*erm* Δ*sipL* with *K35A* in the *spoIVA* locus and *sipL-mCherry* in the *pyrE* locus | This study |
| 2662 | 630Δ*erm* Δ*sipL spoIVA-K35E/ sipL-mCherry* | 630Δ*erm* Δ*sipL* with *K35E* in the *spoIVA* locus and *sipL-mCherry* in the *pyrE* locus | This study |
| 2665 | 630Δ*erm* Δ*sipL spoIVA-T75A/ sipL-mCherry* | 630Δ*erm* Δ*sipL* with *T75A* in the *spoIVA* locus and *sipL-mCherry* in the *pyrE* locus | This study |
| 2668 | 630Δ*erm* Δ*sipL spoIVA-D102A/ sipL-mCherry* | 630Δ*erm* Δ*sipL* with *D102A* in the *spoIVA* locus and *sipL-mCherry* in the *pyrE* locus | This study |
| 2888 | 630Δ*erm* Δ*sipL spoIVA-T75A/ sipL*$_{W475E}$ | 630Δ*erm* Δ*sipL* with *T75A* in the *spoIVA* locus and *sipL*$_{W475E}$ in the *pyrE* locus | This study |
| 2891 | 630Δ*erm* Δ*sipL spoIVA-T75A/ sipL*$_{I463R}$ | 630Δ*erm* Δ*sipL* with *T75A* in the *spoIVA* locus and *sipL*$_{I463R}$ in the *pyrE* locus | This study |
| 2894 | 630Δ*erm* Δ*sipL spoIVA-D102A/ sipL*$_{W475E}$ | 630Δ*erm* Δ*sipL* with *D102A* in the *spoIVA* locus and *sipL*$_{W475E}$ in the *pyrE* locus | This study |
| 2897 | 630Δ*erm* Δ*sipL spoIVA-D102A/ sipL*$_{I463R}$ | 630Δ*erm* Δ*sipL* with *D102A* in the *spoIVA* locus and *sipL*$_{I463R}$ in the *pyrE* locus | This study |
| 2930 | 630Δ*erm* Δ*sipL spoIVA-K35A/ sipL*$_{W475E}$ | 630Δ*erm* Δ*sipL* with *K35A* in the *spoIVA* locus and *sipL*$_{W475E}$ in the *pyrE* locus | This study |
| 2933 | 630Δ*erm* Δ*sipL spoIVA-K35A/ sipL*$_{I463R}$ | 630Δ*erm* Δ*sipL* with *K35A* in the *spoIVA* locus and *sipL*$_{I463R}$ in the *pyrE* locus | This study |

***E. coli* strains**

| Strain # | Strain name | Relevant genotype or features | Source |
|---|---|---|---|
| 41 | DH5α | F– Φ80*lacZ*ΔM15 Δ(*lacZYA-argF*) U169 *recA1 endA1 hsdR17* ($rK^{-}$, $mK^{+}$) *phoA supE44* λ– *thi-1 gyrA96 relA1* | D. Cameron |
| 531 | HB101/pRK24 | F- *mcrB mrr hsdS20*($rB^{-}mB^{-}$) *recA13 leuB6 ara-13 proA2 lavYI galK2 xyl-6 mtl-1 rpsL20* carrying pRK24 | C. Ellermeier |
| 1087 | BTH101 | $F^{-}$, *cya-99, araD139, galE15, galK16, rpsL1, hsdR2, mcrA1, mcrB1* | (11) |
| 1251 | pUT18C-*spoIVA* | pUT18C-*spoIVA* in DH5α | This study |
| 1252 | pKNT25-*sipL* | pKNT25-*sipL* in DH5α | This study |
| 1685 | pMTL-YN1C *spoIVA T75A* | pMTL-YN1C *spoIVA T75A* in HB101 | This study |
| 1686 | pMTL-YN1C *spoIVA D102A* | pMTL-YN1C *spoIVA D102A* in HB101 | This study |
| 1687 | pMTL-YN1C *spoIVA K35A* | pMTL-YN1C *spoIVA K35A* in HB101 | This study |
| 1704 | pMTL-YN3 Δ*sipL* | pMTL-YN3 Δ*sipL* in HB101 | (13) |
| 1710 | pMTL-YN1C *spoIVA K35E* | pMTL-YN1C *spoIVA K35E* in HB101 | This study |
| 1768 | pMTL-YN1C *mCherry-spoIVA* | pMTL-YN1C *mCherry-spoIVA* | (6) |
| 1777 | pMTL-YN1C *sipL-mCherry* | pMTL-YN1C *sipL-mCherry* in HB101 | (13) |
| 1812 | pMTL-YN1C *cotE-mCherry* | pMTL-YN1C *cotE-mCherry* in HB101 | (13) |

| | | | |
|---|---|---|---|
| 1813 | pMTL-YN1C *mCherry-spoIVA*$_{K35E}$ | pMTL-YN1C *mCherry-spoIVA*$_{K35E}$ in HB101 | This study |
| 1818 | pMTL-YN3 *spoIVA K35E* | pMTL-YN3 *spoIVA K35E* in HB101 | This study |
| 1819 | pMTL-YN3 *spoIVA D102A* | pMTL-YN3 *spoIVA D102A* in HB101 | This study |
| 1829 | pMTL-YN1C *mCherry-spoIVA*$_{D102A}$ | pMTL-YN1C *mCherry-spoIVA*$_{D102A}$ in HB101 | This study |
| 1835 | pMTL-YN3 *spoIVA K35A* | pMTL-YN3 *spoIVA K35A* in HB101 | This study |
| 1836 | pMTL-YN3 *spoIVA T75A* | pMTL-YN3 *spoIVA T75A* in HB101 | This study |
| 1837 | pMTL-YN1C *mCherry-spoIVA*$_{K35A}$ | pMTL-YN1C *mCherry-spoIVA*$_{K35A}$ in HB101 | This study |
| 1838 | pMTL-YN1C *mCherry-spoIVA*$_{T75A}$ | pMTL-YN1C *mCherry-spoIVA*$_{T75A}$ in HB101 | This study |
| 1859 | pMTL-YN1C *sipL-3xFLAG* | pMTL-YN1C *sipL-3xFLAG* in HB101 | (14) |
| 1896 | pMTL-YN1C *sipL*$_{\mathrm{I463R}}$ | pMTL-YN1C *sipL*$_{\mathrm{I463R}}$ in HB101 | (14) |
| 1911 | pMTL-YN1C *sipL*$_{\mathrm{W475E}}$ | pMTL-YN1C *sipL*$_{\mathrm{W475E}}$ in HB101 | (14) |
| 2333 | pUT18C-*spoIVA*$_{K35E}$ | pUT18C-*spoIVA*$_{K35E}$ in DH5α | This study |
| 2334 | pUT18C-*spoIVA*$_{D102A}$ | pUT18C-*spoIVA*$_{D102A}$ in DH5α | This study |
| 2359 | pUT18C-*spoIVA*$_{K35A}$ | pUT18C-*spoIVA*$_{K35A}$ in DH5α | This study |
| 2360 | pUT18C-*spoIVA*$_{T75A}$ | pUT18C-*spoIVA*$_{T75A}$ in DH5α | This study |

**Plasmids**

| | | |
|---|---|---|
| pMTL-YN1C | For cloning complementation constructs to be integrated into the *pyrE* locus of 630Δ*erm*Δ*pyrE* | (15) |
| pMTL-YN3 | For cloning allelic exchange constructs to modify 630Δ*erm*Δ*pyrE* | (15) |
| pUT18C | For cloning *spoIVA* as a C-terminal fusion to the T18 fragment from adenylate cyclase to *spoIVA* | (11) |
| pKNT25 | For cloning *sipL* as an N-terminal fusion to the T25 fragment from adenylate cyclase to *sipL* | (11) |

### Table S2. Primers used in this study.

| Number | Primer name | |
|---|---|---|
| 711 | 5' *spoIVA* K35E SOE | GTTGGACCTGTAAGAACAGGAGAATCAACTTTTATAAGAAAATTTATGGAAAAGTTGG |
| 712 | 3' *spoIVA* K35E rev oes | CCAACTTTTCCATAAATTTTCTTATAAAAGTTGATTCTCCTGTTCTTACAGGTCCAAC |
| 1215 | 5' *spoIVA* T75A PCR SOE | GGTAAAACGATAATGGCAGTAGAACCAAAATTTG |
| 1216 | 3' *spoIVA* T75A PCR rev OES | CAAATTTTGGTTCTACTGCCATTATCGTTTTACC |
| 1217 | 5' *spoIVA* D102A PCR SOE | GTAAGAATGGTAGCTTGTGTTGGATACATAG |
| 1218 | 3' *spoIVA* D102A PCR rev OES | CTATGTATCCAACACAAGCTACCATTCTTAC |
| 1337 | 3’ KpnI *spoIVA* | AAAGGTACCTTATAACAAAATAGTTATAATATTAG |
| 1453 | 5' BamHI *sipL* | AAAGGATCCAATGGAATTAATTAAAGATGTAATTAAAG |
| 1454 | 3' KpnI *sipL* | AAAGGTACCAAATCTACTAATACGAC |
| 1452 | 5’ BamHI *spoIVA* | AAAGGATCCAATGTATAGGAGGAATAATATGAATAATAAC |
| 1482 | 5' *spoIVA* K35A SOE | GGACCTGTAAGAACAGGAGCATCAACTTTTATAAGAAAATTTATGGAAAAGTTGG |
| 1483 | 3' *spoIVA* K35A SOE rev | CCAACTTTTCCATAAATTTTCTTATAAAAGTTGATGCTCCTGTTCTTACAGGTCC |
| 1927 | 5' AscI Δ*spoIVA* 1043 | AACGGCGCGCCCCATTTATAAAAGAAGGACAAGTTATTG |
| 1928 | 3' SbfI Δ*spoIVA* 975 bp | ATTATTCCTGCAGGACCTTTTATTTCATCACTAAATACACC |
| 2036 | 5' NotI *spoIVA* gibson | TTAGGGATGTAATAAGCGGCCGCCAATTAGCATTGTAGTTTACTAGTTTTTGTATATAGG |
| 2037 | 3' XhoI *spoIVA* gibson | CAAGCTTGCATGTCTGCAGGCCTCGAGCTTTGAAACAATCCTGTCGAAACATATAC |
| 2133 | 3' XhoI mCherry Gibson | GCCAAGCTTGCATGTCTGCAGGCCTCGAGTTATTTATATAATTCATCCATACCTCCTGTTG |
| 2202 | 5' $P_{spoIVA}$-mCherry SOE | CCCTTAACTCTAAATATAATTTAATAAGATTAGATGGTATCTAAAGGAGAAGAAGATAAT |
| 2203 | 3' $P_{spoIVA}$-mCherry rev eos | ATTATCTTCTTCTCCTTTAGATACCATCTAATCTTATTAAATTATATTTAGAGTTAAGGG |
| 3064 | 5' BamHI *spoIVA* T18 | CACTGCAGGTCGACTCTAGAGGATCCCATGTATAGGAGGAATAATATGAATAATAAC |
| 3065 | 3' KpnI *spoIVA* T18 | GTGCACCATATTACTTAGTTATATCGATGAATTCGAGCTCGGTACCTTATAACAAAATAG |

Restriction sites are underlined.

### References


1. Bennison DJ, Irving SE, Corrigan RM. 2019. The Impact of the Stringent Response on TRAFAC GTPases and Prokaryotic Ribosome Assembly. Cells 8.
2. Ramamurthi KS, Clapham KR, Losick R. 2006. Peptide anchoring spore coat assembly to the outer forespore membrane in *Bacillus subtilis*. Mol Microbiol 62:1547-1557.
3. Castaing JP, Lee S, Anantharaman V, Ravilious GE, Aravind L, Ramamurthi KS. 2014. An autoinhibitory conformation of the *Bacillus subtilis* spore coat protein SpoIVA prevents its premature ATP-independent aggregation. FEMS Microbiol Lett 358:145-53.
4. Castaing J-P, Nagy A, Anantharaman V, Aravind L, Ramamurthi K. 2013. ATP hydrolysis by a domain related to translation factor GTPases drives polymerization of a static bacterial morphogenetic protein. Proc Natl Acad Sci U S A 110:60.
5. Fimlaid KA, Jensen O, Donnelly ML, Siegrist MS, Shen A. 2015. Regulation of *Clostridium difficile* Spore Formation by the SpoIIQ and SpoIIIA Proteins. PLoS Genet 11:e1005562.
6. Ribis JW, Ravichandran P, Putnam EE, Pishdadian K, Shen A. 2017. The Conserved Spore Coat Protein SpoVM Is Largely Dispensable in *Clostridium difficile* Spore Formation. mSphere 2.
7. Ebmeier SE, Tan IS, Clapham KR, Ramamurthi KS. 2012. Small proteins link coat and cortex assembly during sporulation in *Bacillus subtilis*. Mol Microbiol 84:682-96.
8. Fimlaid KA, Bond JP, Schutz KC, Putnam EE, Leung JM, Lawley TD, Shen A. 2013. Global Analysis of the Sporulation Pathway of *Clostridium difficile*. PLoS Genet 9:e1003660.
9. Degasperi A, Birtwistle MR, Volinsky N, Rauch J, Kolch W, Kholodenko BN. 2014. Evaluating strategies to normalise biological replicates of Western blot data. PLoS One 9:e87293.
10. Horton R, Hunt H, Ho S, Pullen J, Pease L. 1989. Engineering hybrid genes without the use of restriction enzymes: gene splicing by overlap extension. Gene 77:61-68.
11. Karimova G, Pidoux J, Ullmann A, Ladant D. 1998. A bacterial two-hybrid system based on a reconstituted signal transduction pathway. Proc Natl Acad Sci U S A 95:5752-6.
12. Donnelly ML, Li W, Li YQ, Hinkel L, Setlow P, Shen A. 2017. A *Clostridium difficile*-Specific, Gel-Forming Protein Required for Optimal Spore Germination. MBio 8.
13. Ribis JW, Fimlaid KA, Shen A. 2018. Differential requirements for conserved peptidoglycan remodeling enzymes during *Clostridioides difficile* spore formation. Mol Microbiol 110:370-389.
14. Touchette MH, Benito de la Puebla H, Ravichandran P, Shen A. 2019. SpoIVA-SipL Complex Formation Is Essential for *Clostridioides difficile* Spore Assembly. J Bacteriol 201.
15. Ng YK, Ehsaan M, Philip S, Collery MM, Janoir C, Collignon A, Cartman ST, Minton NP. 2013. Expanding the repertoire of gene tools for precise manipulation of the *Clostridium difficile* genome: allelic exchange using *pyrE* alleles. PLoS One 8:e56051.
